# The Habenular Receptor GPR151 Regulates Addiction Vulnerability Across Drug Classes

**DOI:** 10.1101/720508

**Authors:** Beatriz Antolin-Fontes, Kun Li, Jessica L. Ables, Michael H. Riad, Andreas Görlich, Maya Williams, Cuidong Wang, Sylvia M. Lipford, Maria Dao, Henrik Molina, Jianxi Liu, Paul J. Kenny, Ines Ibañez-Tallon

## Abstract

The habenula controls the addictive properties of nicotine but also densely expresses opioid and cannabinoid receptors. As such, identification of strategies to manipulate habenular activity may yield new approaches to treat substance use disorders. Here we show that GPR151, an orphan G protein-coupled receptor (GPCR) highly enriched in the habenula of humans and rodents plays a critical role in regulating habenular function and behavioral responses to addictive drugs. We show that GPR151 is expressed on axonal and presynaptic membranes and synaptic vesicles, and regulates synaptic fidelity and plasticity. We find that GPR151 associates with synaptic components controlling vesicle release and ion transport and couples to the G-alpha inhibitory protein Gα_o1_ to reduce cAMP levels. Stable cell lines expressing GPR151 confirm that it signals via Gi/o and are amenable to ligand screens. *Gpr151* null mice show diminished behavioral responses to nicotine, and self-administer greater quantities of the drug, phenotypes rescued by viral re-expression of *Gpr151* in the habenula. *Gpr151* null mice are also insensitive to the behavioral actions of morphine and cannabinoids. These data identify GPR151 as a critical modulator of habenular function that controls addiction vulnerability across different drug classes.

**Highlights:**

- Habenula neurons are enriched in nicotinic, opioid, cannabinoid and GPR151 receptors
- GPR151 modulates synaptic fidelity and release probability at habenular terminals.
- Habenular GPR151 plays a role in drug abuse and food intake/weight control
- GPR151 couples to the G-alpha inhibitory protein Gα_o1_ to reduce cAMP levels.

**eTOC Blurb:** Antolin-Fontes at al. identify a G protein-coupled receptor, GPR151, which is highly enriched in human habenular neurons. These neurons are primarily enriched with nicotinic, opioid and cannabinoid receptors. We find that GPR151 modulates habenular synaptic vesicle release probability and behavioral responses to these drugs of abuse.

## INTRODUCTION

A fundamental role for the habenula in the control of impulsive or compulsive behaviors, and in the devaluation of reward by delay and effort, has been supported strongly by several recent studies (Kobayashi et al., 2013; Lammel et al., 2012; Proulx et al., 2014). The conservation of this ancient structure across vertebrate evolution, and the many behavioral abnormalities evident in animals carrying habenular lesions (Agetsuma et al., 2010; Kobayashi et al., 2013), further support the crucial roles of the habenula in processing reward-related and aversive signals. However, it has become evident that alterations in the functions of this pathway are also associated with psychiatric disorders including obsessive compulsive disorder (Proulx et al., 2014) and drug addiction (Antolin-Fontes et al., 2015; McLaughlin et al., 2017).

The global impact of substance abuse and addiction disorders on health and economy is well documented. Currently, about 1.3 billion people consume tobacco and 185 million people use psychoactive drugs (WHO), while the Department of Health and Human services estimates that 2.1 million people in the US have an opioid use disorder, while another 11.2 million misuse opioids. Irrespective of the type of drug, addiction is a chronic relapsing disorder characterized by compulsive drug seeking, escalation of intake, and development of affective and physical symptoms of withdrawal upon abrupt discontinuation or decrease in intake (Koob and Volkow, 2016). Animal studies and human imaging analyses have revealed that many psychoactive drugs act on the mesocorticolimbic dopamine system, a circuit that appears to be common to the rewarding effects of some drugs of abuse, as well as other reinforcing natural behaviors such as eating, thirst, gambling and sexual drive (Lammel et al., 2014; Lammel et al., 2012; Nutt et al., 2015).However recent findings challenge the view that DA release has an exclusive role in reward processing and that stress, and aversive stimuli activate and remodel partially overlapping networks within the mesocorticolimbic dopamine (DA) system (Ostroumov and Dani, 2018). More recently, other structures have been implicated in reward and aversion, including the habenula (Antolin-Fontes et al., 2015; Boulos et al., 2019; Mathis and Kenny, 2018). Increasing evidence suggests that the degree of sensitivity to both the rewarding and aversive aspects of an addictive drug and the severity of the withdrawal after discontinuing its use contribute to the addiction process (Koob and Volkow, 2016). Understanding the mechanisms of these processes may reveal novel insights into the mechanics of the disorder and identify new targets for medications development.

The medial habenula (MHb) and its major projection site, the interpeduncular nucleus (IPN) are highly enriched in nicotinic acetylcholine receptors (nAChRs) (Ables et al., 2017; Changeux, 2010; Gorlich et al., 2013; Shih et al., 2014) and are key in the control of nicotine intake, aversion, withdrawal and relapse (Fowler et al., 2011; Frahm et al., 2011; Gorlich et al., 2013; Salas et al., 2009). The MHb also densely expresses mu-opioid (MOR) and cannabinoid 1 (CB1R) receptors. It has been implicated in morphine dependence and withdrawal (Neugebauer et al., 2013), aversion to MOR blockade (Boulos et al., 2019), aversive memories mediated by CB1R (Soria-Gomez et al., 2015) and depression induced by decreased habenular CB1R during nicotine abstinence (Simonnet et al., 2017). Furthermore, it has been shown that chronic exposure to D-amphetamine, methamphetamine, MDMA, cocaine, or nicotine can induce degeneration of the fasciculus retroflexus (FR), the main output tract of the habenula (Velasquez et al., 2014). The finding that this descending pathway is compromised following drug binges has implications not only for theories of drug addiction but also for psychosis in general (Ellison, 2002). Thus, the emerging picture is that the Hb-IPN pathway acts as an inhibitory motivational signal that limits drug intake (Fowler et al., 2011), and that alterations in the functioning of this pathway by drug consumption may contribute to many aspects of addiction.

Here we investigated GPR151, an orphan GPCR with remarkably selective expression in habenular axonal projections. We found that GPR151 is highly conserved in the human Hb-IPN circuit and specifically localizes to axonal and presynaptic membranes and synaptic vesicles (SV). GPR151 partially overlaps with α3β4-containing nAChRs, MOR and CB1R receptors at habenular terminals in the IPN. We observed reduced behavioral responses to drugs that act at these receptors in *Gpr151*-knockout (KO) mice. Furthermore, *Gpr151*-KO mice intravenously self-administered much higher amounts of nicotine than their wildtype (WT) counterparts, consistent with a role for GPR151 in regulating the stimulatory effects of nicotine on habenular aversion circuits. Mass spectrometry identified SV proteins, presynaptic ATPases and the G-proteins Gαo1 and Gβ1 as GPR151 interacting proteins, indicating that GPR151 couples to the G-alpha inhibitory pathway to reduce cAMP. In the absence of a ligand we employed optogenetics to stimulate MHb terminals and found increased synaptic fidelity rates and smaller evoked EPSCs in *Gpr151*-KO mice. Taken together, our data reveal a key role for GPR151, and the MHb-IPN circuit more broadly, in regulating the sensitivity and aversion to addictive drugs across class, and suggest that small molecule modulators of GPR151 activity may represent an entirely new class of addiction therapeutics.

## RESULTS

### GPR151 is specifically co-expressed in the habenula-IPN pathway with nicotinic, opioid and cannabinoid receptors

GPR151 expression is remarkably conserved and restricted to the habenula-IPN tract in the brains of zebrafish, mice and rats (Broms et al., 2015). Most GPR151-expressing cells are concentrated in the Hb (mostly in the MHb and few scattered cells in the LHb (Broms et al., 2015; Kobayashi et al., 2013). Notably, GPR151 is not present in the soma of habenular neurons, but along their axonal projections that comprise the FR and at terminals in the IPN (Figure 1A-D). To investigate whether the pattern of GPR151 expression was also conserved in humans, we obtained post-mortem adult human brain samples containing part of the diencephalon and midbrain (Figure 1E-O). Immunostaining analyses showed that GPR151 specifically labeled habenular projections arising in the MHb and LHb (Figure 1G-I). We detected very strong GPR151 signal along the FR in horizontal human brain sections (Figure 1J-L), as well as in the IPN (Figure 1M-O) where MHb projections terminate. The specific immunoreactivity of human Hb and IPN to GPR151 antibody is identical to the pattern we observed in rodents and zebrafish (Broms et al., 2015). Western blot analyses in GPR151 transfected cells, WT mice and human IPN samples revealed two bands corresponding to GPR151: one of the expected size at 46 kDa and an additional higher molecular band (≈53 kDa) suggesting posttranslational modifications (Figure 1P). These results show that GPR151 is specifically expressed in the Hb-IPN axonal tract in humans, suggesting that its function is likely to be conserved across vertebrates.

**Figure 1:**
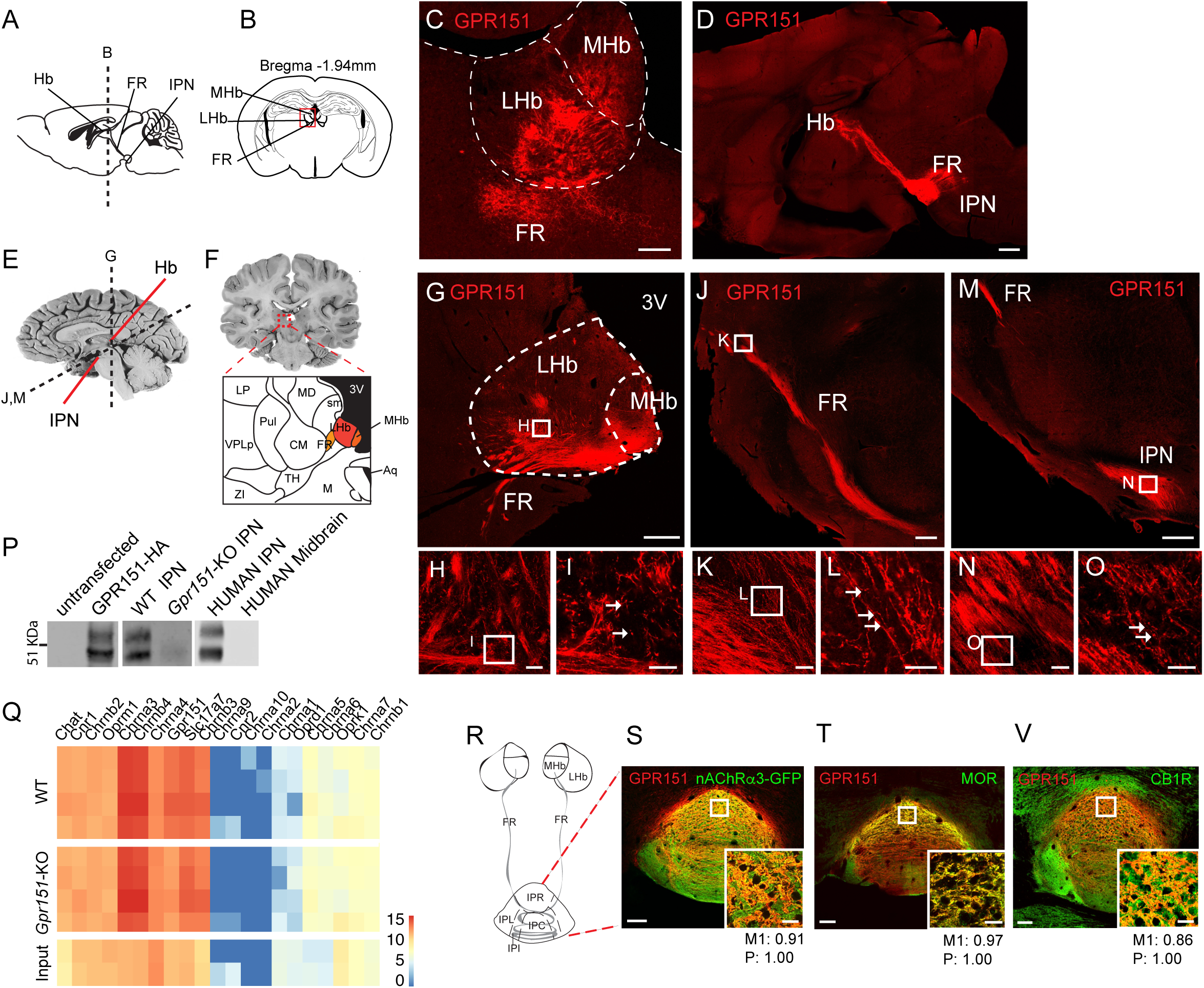
The habenula-IPN pathway is enriched in GPR151, nicotinic, opioid and cannabinoid receptors. (A-B) Sagittal and coronal mouse brain scheme indicating the habenula (medial (MHb) and lateral (LHb), fasciculus retroflexus (FR) and interpeduncular nucleus (IPN). (C-D) GPR151 is expressed in the axonal fibers originating mostly in MHb and passing through the LHb to form the FR (C; Scale bar: 100 µm); and along the FR and at axonal terminals in the IPN (D; Scale bar: 500 µm). (E-F) Sagittal and coronal human brain sections indicating the habenula and IPN. Dotted lines show the plane of section in G,J, and M. Image modified from (Haines, 2004). Diagram highlighting the MHb, LHb and FR (adapted from Allen Brain Human Atlas). (G-I) Coronal and horizontal sections of human brain samples showing GPR151 immunostaining in the MHb, LHb and beginning of the FR (G), along the FR (J) and IPN (M). Punctuate pattern of GPR151 was observed along the axons and in the IPN (arrows) on high magnification pictures of the square areas indicated. Scale bar: 1mm in G, 50) µm in J and M, 50 µm in H, K and N and 20 µm in I, L and O. (P) Western blot analysis of untransfected and GPR151-HA transfected HEK293 cells; immunoprecipitated IPN samples from WT, *Gpr151*-KO mice, human IPN and human midbrain. Two bands (46 and 53 kDa) correspond to GPR151. (Q) TRAP data collected from MHb neurons of *Chat-EGFP-L10a* mice crossed to WT and *Gpr151*-KO show high and similar expression levels (z-score transformed normalized counts) of *Chat*, opioid (*Oprm1*) cannabinoid *(Cnr1*) and nicotinic (*Chrnb2,Chrna3 Chrnb4, Chrna4, Chrnb3*) and *Slc17a7* (VGLUT1) translated mRNAs in IP samples compared to Input samples. Only *Gpr151* shows differential expression between WT and *Gpr151-KO* IP samples (n=6 WT, 6 KO). (R) Scheme of the Hb-IPN subnuclei (IPR: rostral, IPL: lateral, IPI: intermediate, IPC: central). (S-V) Double immunostaining of GPR151 and nAChRα3-GFP (S), µ-opioid receptor (MOR) (T) and cannabinoid receptor 1 (CB1R) (V) showing high coexpression in the IPR where Manders’ co-localization index (M1) and significance of correlation (P) were measured (indicated square area). Scale bar: 100 µm for low magnification pictures, 20 µm for high magnification pictures.

To obtain a quantitative list of highly translated mRNAs corresponding to receptors expressed in the same neurons as *Gpr151*, we employed *Chat-EGFP-L10a* transgenic mice for Translating Ribosome Affinity Purification (TRAP) analysis (Doyle et al., 2008). Six biological replicates were collected from the habenulae of *Chat-EGFP-L10a* mice and the resulting INPUT and Immunoprecipitated (IP) mRNAs were analyzed by RNA-Seq (Figure 1Q and Table S1). The high enrichment of *Gpr151* in the TRAP IP (Figure S1A, S1B, S2A and Table S1) is consistent with its expression in habenular cholinergic neurons, and indicates that other receptors co-enriched in the IP fraction are also enriched in habenular neurons. These analyses demonstrate high co-enrichment of *Gpr151* with *Chrna3* and *Chrnb4*, and significant overlap with *Oprm1* and *Cnr1* mRNAs in the *Chat* MHb population (Figure 1Q and S2A).

To confirm the TRAP results, we performed immunofluorescence analyses in *Chrna3/eGFP* mice, which label α3β4 nAChR positive neurons in the brain (Frahm et al., 2011). Intense nAChRα3-eGFP expression was evident in MHb neurons and their projections to the IPN (Figure S3B, S3E) and demonstrated considerable overlap with GPR151 (co-localization index: 0.91) in the central and rostral part of the IPN (IPC and IPR) (Figure 1S and Figure S3E-F). Similarly, co-immunostaining of GPR151 with the µ-opioid receptor (MOR) antibody showed intense colocalization in axonal terminals in the IPR (colocalization index: 0.97) (Figure 1T and Figure S3G-H) and partial overlap in the lateral part of the MHb (Figure S3C).

Double immunostaining of GPR151 and the cannabinoid receptor 1 (CB1R) revealed intense costaining in habenular axons terminals in the IPR (colocalization index: 0.86) (Figure 1V and Figure S3I-J) and low expression in habenular cell bodies (Figure S3D). Single stains of MOR and CB1R (Figure S3) are consistent with their reported patterns in habenula and IPN (Gardon et al., 2014; Soria-Gomez et al., 2015). Importantly, gene deletion of *Gpr151* in mice (*Gpr151*-KO mice) does not influence the levels of translated mRNAs of *Oprm1* and *Cnr1* in MHb neurons quantified by TRAP (Figure 1Q), nor the expression of MOR and CB1R at habenular terminals (Figure S4), indicating that GPR151 expression does not modulate expression of these receptors localized at the same habenular terminals nor induce adaptive responses in their expression in the mutant mice. Together, the TRAP and IHC results establish that GPR151 is highly enriched in habenular projections in the IPN that also express α3β4 nAChRs, MOR and CB1R, and raise the possibility that GPR151 may influence the actions of of nicotine, morphine and cannabinoids in the habenula and thereby regulate behavioral responses to these drugs.

### Reduced sensitivity to nicotine, morphine and the cannabinoid agonist ACEA in *Gpr151*-KO mice

Given the high-degree of conservation of GPR151 between rodent and human brains and the co-expression of GPR151 with receptors for multiple drugs of abuse, we examined the behavioral responses in mice lacking this receptor. This is important because the findings could be potentially relevant to human disorders with reward-related abnormalities, such as affective and addictive disorders. We first tested *Gpr151*-KO mice at baseline conditions and found no differences in locomotor activity, anxiety-like behaviors measured by the elevated plus maze, sensorimotor gating analyzed by the prepulse inhibition test, and anhedonic-like behavior measured by the preference of sucrose consumed in a two-bottle choice experiment (Figure S5A-D), suggesting that GPR151 does not regulate affective-related behaviors at baseline.

We next examined the behavioral responses of *Gpr151*-KO mice to nicotine, morphine and a synthetic cannabinoid agonist. We assayed locomotor activity after an acute nicotine challenge, which reflects the sensitivity of an individual to nicotine (Clarke and Kumar, 1983). Baseline activity of *Gpr151*-KO mice was similar to WT (min 0 in Figure 2A, day 0 in Figure 2B, saline in Figure 2C and Figure S5A). However, acute nicotine-induced hypolocomotion was significantly less prominent in *Gpr151*-KO mice (min 5, 10 in Figure 2A, day 1 in Figure 2B), indicating that *Gpr151*-KO mice have reduced sensitivity to the motor-depressing property of nicotine. While both WT and KO mice displayed similar nicotine-induced hypolocomotion at the 0.35 mg/kg dose, unlike the WT mice, *Gpr151*-KO hypolocomotion was not further increased at 0.65 and 1.5 mg/kg nicotine (Figure 2C). In addition, daily injections of nicotine induced tolerance to the hypolocomotor effect of nicotine in WT mice after 9 days of treatment, but *Gpr151*-KO mice did not demonstrate an adaptive response to repeated injection of nicotine (Figure 2B, day 9-11). These results show that *Gpr151*-KO mice are less sensitive to high doses of nicotine and do not show behavioral plasticity in response to repeated exposures of nicotine.

**Figure 2.**
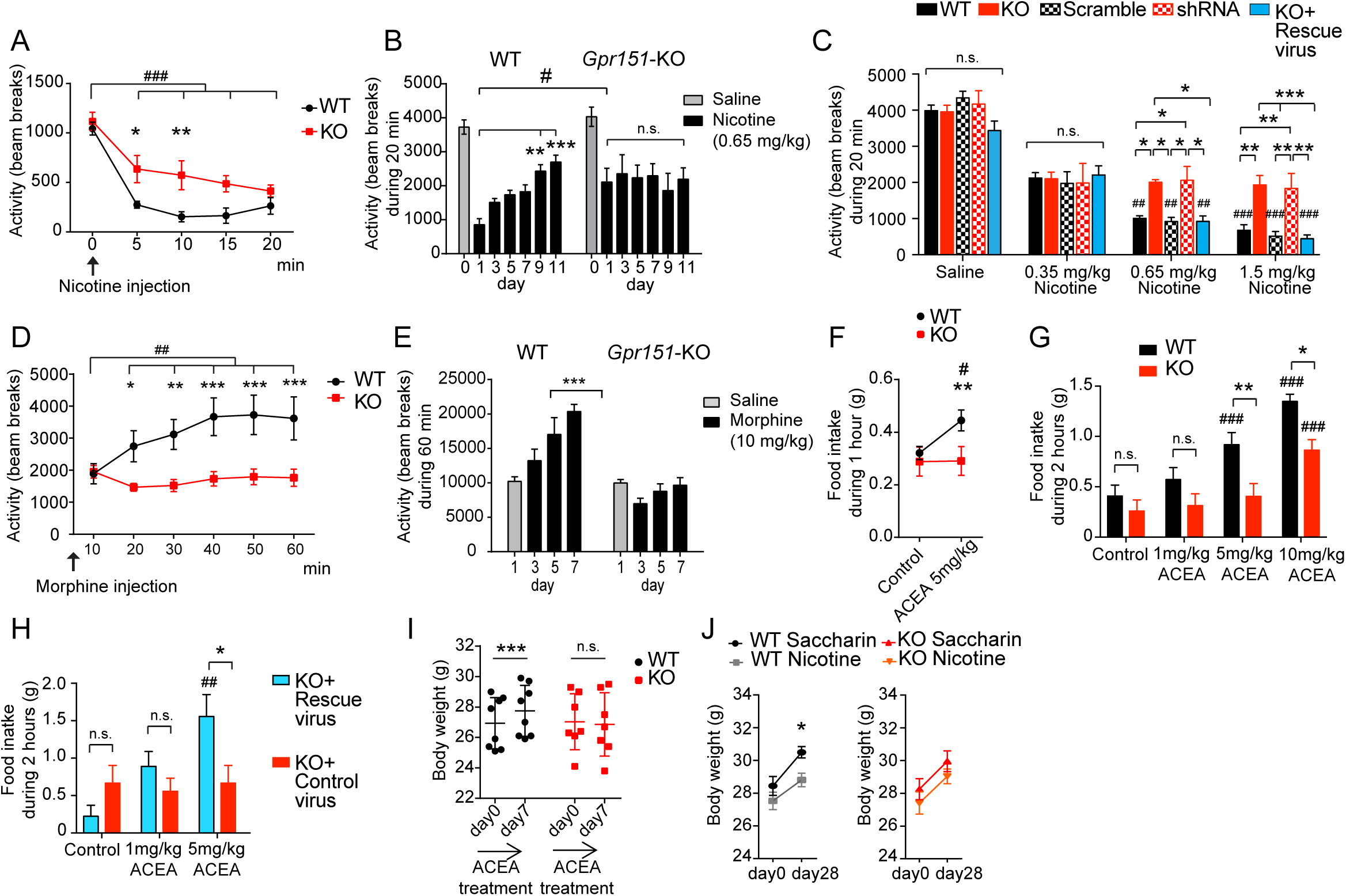
*Gpr151*-KO mice show blunted behavioral responses to nicotine, morphine and cannabinoids. (A) Hypolocomotion induced by acute injection of nicotine (0.65Lmg/kg, i.p.), measured as number of beam breaks in 5 min intervals, is significantly diminished in *Gpr151*-KO mice (n=7 per genotype; RM 2-way ANOVA, Bonferroni multiple comparisons *p<0.05, **p<0.01 for genotype; Tukey’s multiple comparisons test ###p<0.001 for time in both groups). Basal locomotor activity (min 0) is similar between groups. (B) Consecutive daily injections of nicotine induce tolerance to the hypolocomotor effects of nicotine in WT but not in *Gpr151*-KO mice (n=7 per genotype, RM 2-way ANOVA, Bonferroni multiple comparisons #p<0.05 for genotype; Tukey’s multiple comparisons test **p<0.01, ***p<0.001 for time). (C) Hypolocomotion induced by nicotine is observed at the higher 2 doses of nicotine (0.65 and 1.5 mg/kg) in WT mice, WT mice injected with AAV2.1-scramble-shRNA (scramble) and in *Gpr151*-KO mice injected with AAV2.1-CAG-*Gpr151* (rescue) in the MHb, but not in *Gpr151*-KO mice and in WT mice injected with AAV2.1-shRNA against *Gpr151* (shRNA) in the MHb. No differences were observed between genotypes upon saline injection or a low dose of nicotine (0.35mg/kg). (n=7-8 per genotype, RM 2-way ANOVA, Tukeýs multiple comparison test *p<0.05, **p<0.01, ***p<0.001 for genotype; ##p<0.01, ###p<0.001 compared to 0.35mg/kg nicotine within the same genotype). (D) Acute injection of morphine (10Lmg/kg, s.c.) induces hyperlocomotion in WT mice, but not in *Gpr151*-KO mice (n=7-8 per genotype; RM 2-way ANOVA, Bonferroni multiple comparisons *p<0.05, **p<0.01, ***p<0.001 for genotype; Tukey’s multiple comparisons test ##p<0.01 for time). (E) Consecutive daily injections of morphine induce sensitization to hyperlocomotor activity in WT mice but not in *Gpr151*-KO mice. (n=7-8 per genotype; RM 2-way ANOVA, Tukey’s multiple comparisons test ***p<0.001). (F) Acute injection of the cannabinoid agonist ACEA (5mg/kg, i.p.) increases food intake in WT mice but not in *Gpr151*-KO mice (n=8-7 per genotype; RM 2-way ANOVA, Bonferroni’s multiple comparisons test, **p<0.01 in WT mice when comparing saline injection to ACEA injection, #p<0.05 when comparing WT to *Gpr151*-KO injected with ACEA). (G) Hyperphagia induced by ACEA is observed at the higher 2 doses of ACEA (5 and 10 mg/kg) in WT mice and only at the high dose in *Gpr151*-KO mice (n=10-8 per genotype; RM 2-way ANOVA, Bonferroni’s multiple comparisons test for genotype, *p<0.05, **p<0.01; Tukey’s multiple comparisons test for ACEA concentration compared to saline, ###p<0.0001). (H) *Gpr151*-KO mice injected with AAV2.1-CAG-*Gpr151* in the MHb (rescue virus) but not with control virus show increased food intake after injection of ACEA (5mg/kg) (n=9 per genotype; RM 2-way ANOVA, Bonferroni’s multiple comparisons test for genotype, *p<0.05; Tukey’s multiple comparisons test for ACEA concentration compared to saline, ## p<0.01). (I) Daily injections of ACEA (5mg/kg, i.p.) for 7 days increase the body weight of WT mice but not of *Gpr151*-KO mice (n=8-7 per genotype; RM 2-way ANOVA, Bonferroni’s multiple comparisons test, ***p<0.001). (J) Mice consuming 162,5 µg/ml nicotine + 2% saccharin in the drinking water for 28 days show less increase of body weight with respect to mice treated with only 2% saccharin. No effect of nicotine on body weight was observed between *Gpr151*-KO mice treated with saccharin or nicotine (n=11-26 per group; RM 2-way ANOVA, Sidak’s multiple comparisons test, *p<0.05 when comparing WT saccharin to WT nicotine on day 28). Data are represented as mean ± SEM. See Table S3 for details of statistical analysis.

Next, to rule out a developmental effect caused by the lack of *Gpr151* as the reason for the altered sensitivity to nicotine observed in *Gpr151*-KO mice, we knocked downed and rescued *Gpr151* in the habenula of WT and *Gpr151*-KO mice, respectively. We tested several different shRNAs against *Gpr151* in HEK293T cells and determined that shRNA-3 successfully reduced GPR151 expression by Western blot (Figure S6A-B), and we chose it for *in vivo* analysis. Further, we determined that AAV2/1 yielded the highest level of expression in MHb neurons compared to other serotypes, and generated AAVs containing the shRNA or a scramble-shRNA as a control to knock-down *Gpr151* in WT mice and an AAV containing CAG-*Gpr151* for the rescue experiments in *Gpr151*-KO mice. Viruses were injected bilaterally into the MHb and presence or absence of GPR151 expression in the habenula, along the FR and at the IPN was validated in the injected mice by immunohistochemistry (Figure S6C). The absence of nicotine-induced hypolocomotor effects observed in *Gpr151*-KO mice was recapitulated in shRNA-injected mice while re-expression of *Gpr151* in the KO restored sensitivity to nicotine-induced hypolocomotion (Figure 2C), indicating that GPR151 acts in the MHb to regulate sensitivity to nicotine in adult mice.

Given that *Gpr151-*KO mice have an altered response to nicotine and that GPR151 is co-expressed with MOR and CB1R, we next evaluated the behavioral response of *Gpr151*-KO mice to morphine and the cannabinoid agonist arachidonyl-2′-chloroethylamide (ACEA). As expected, an acute injection of morphine resulted in progressive increase of locomotor activity in WT mice (Figure 2D) (Brase et al., 1977). In contrast *Gpr151*-KO mice were insensitive to the acute hyperlocomotor effect of morphine (Figure 2D). We monitored the development of sensitization to morphine over the course of 6 days of daily injections of morphine. WT mice developed locomotor sensitization while *Gpr151*-KO mice were resistant to sensitization (Figure 2E). Injection of the CB1R agonist ACEA induced hypoactivity as previously described (Almeida et al., 2014), which was indistinguishable between WT and *Gpr151*-KO mice (Figure S5E-F) consistent with studies showing that habenular CB1R does not influence locomotion (Soria-Gomez et al., 2015). However, since CB1R is critical for central regulation of food intake (Koch et al., 2015), we tested whether activation of CB1R induced a differential feeding response in *Gpr151*-KO mice. An injection of ACEA increased feeding in WT mice as described (Koch et al., 2015) while *Gpr151*-KO mice were insensitive to the hyperphagic effect of ACEA at 5mg/kg, and the hyperphagic effect at 10mg/kg was significantly lower than WT mice (Figure 2F-G and Figure S5G-H). Similar to WT mice, *Gpr151*-KO mice injected in the MHb with the rescue virus demonstrated significantly increased food intake upon a 5mg/kg injection of ACEA (Figure 2H). Consecutive injections of ACEA over a 7-day period resulted in significant body weight gain in WT mice but not in *Gpr151*-KO mice (Figure 2I). Body weight of WT and *Gpr151*-KO mice on regular water and food was similar across age (Figure S5I). These data revealed an unsuspected role of GPR151 in feeding behavior induced by activation of CB1R. Prompted by this finding, and given the well-known anorectic effects of nicotine, we tested whether activation of nAChRs resulted in differential weight changes in *Gpr151*-KO mice. Mice were given drinking water containing nicotine and saccharin (to mask the bitter taste of nicotine) for 4 weeks. Control WT mice drinking only saccharin gained significantly more weight than WT mice treated with nicotine (Figure 2J). However, no differences in body weight were observed between *Gpr151*-KO mice treated either with saccharin or nicotine, indicating that *Gpr151*-KO mice are insensitive to the anorectic effects of nicotine.

These data suggest that GPR151 affects the sensitivity to the locomotor effects, feeding patterns and weight gain changes induced by nicotine, opioids and cannabinoids. The demonstration that *Gpr151*-KO mice do not differ from WT mice under baseline conditions, but respond differently to nicotine, morphine and ACEA, supports a critical role for GPR151 signalling in the MHb in controlling behavioral response to drugs of abuse across different classes. In addition, these results reveal an unsuspected role of the habenula in feeding regulation and weight control/energy balance in response to drugs of abuse.

### Reduced nicotine-induced suppression of feeding and increased self-administration of high nicotine doses in *Gpr151*-KO mice

To investigate whether GPR151 controls the reinforcing properties of addictive drugs, we examined WT and *Gpr151*-KO mice in a self-administration task. We focused our attention on nicotine because the MHb-IPN pathway is well-known to control consumption of this drug. Mice first underwent training to respond for food rewards in operant chambers where presses on the active lever resulted in the delivery of food pellets under a fixed-ratio 5 time-out 20 sec (FR5TO20) schedule of reinforcement. No differences were observed in the acquisition of lever pressing behavior (Figure S7). Based on the findings described above, suggesting that the MHb regulates the anorectic actions of nicotine and other drugs of abuse, we determined if nicotine (1mg/kg, s.c.) can reduce responding for food rewards in this task in WT and *Gpr151*-KO mice. In saline treated mice, the number of food pellets earned by WT mice (9.9 +/-0.8) and *Gpr151*-KO mice (11.8 +/-0.5) was comparable (Figure 3A). However, upon nicotine administration WT mice responded significantly less for food rewards than saline WT controls, consistent with the anorectic effect of nicotine on food intake. Similar to our findings with nicotine in the drinking water, this anorectic response was much less pronounced in *Gpr151*-KO mice (Figure 3A) reflecting a reduced sensitivity to nicotine-induced suppression of appetite.

**Figure 3.**
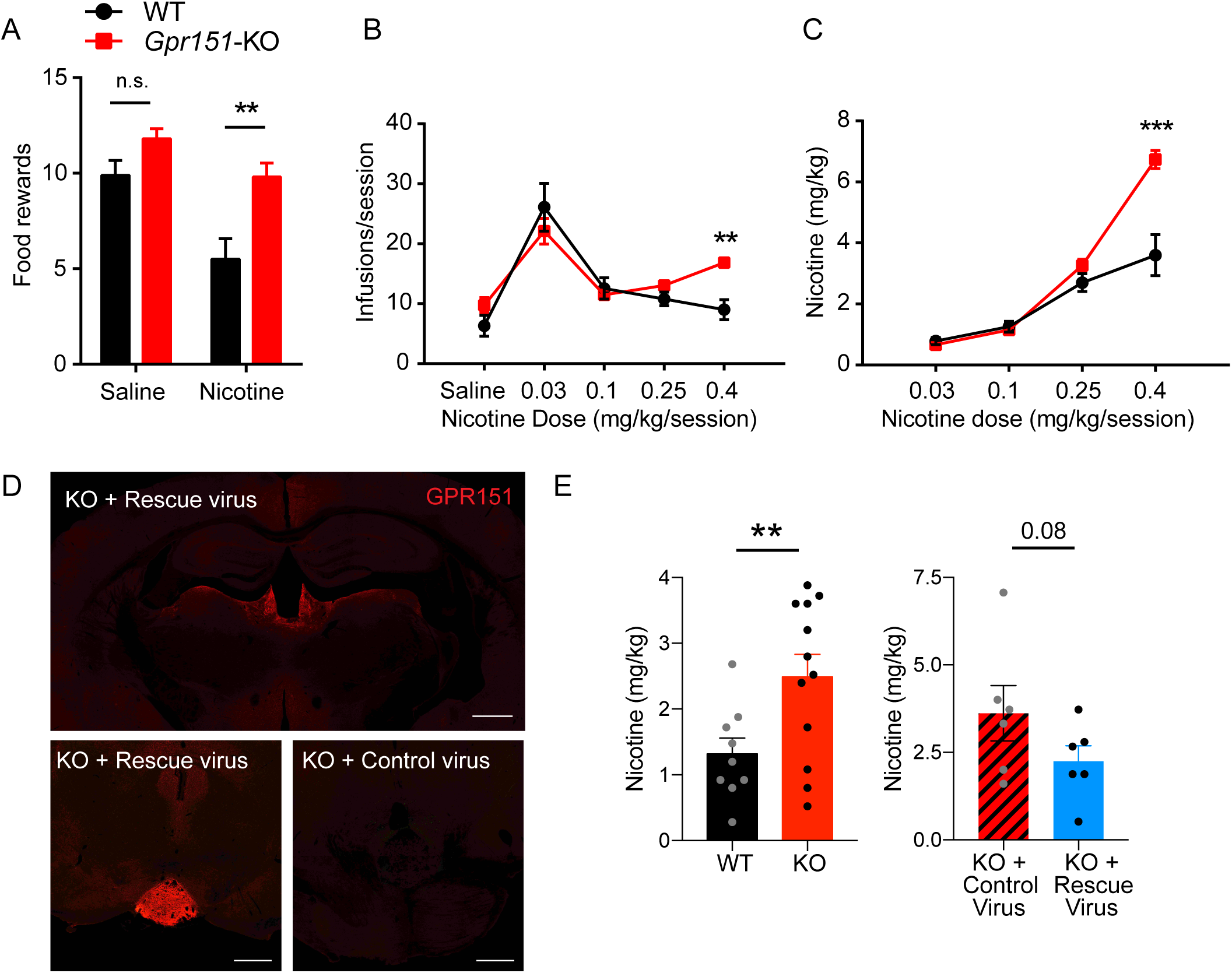
*Gpr151*-KO mice self-administer more nicotine at a high dose. (A) Average number of food rewards after injection of saline or nicotine (1mg/kg, s.c.). *Gpr151*-KO mice are resistant to the anorectic effect of nicotine (n=10 per group, RM 2-way ANOVA, Bonferroni’s multiple comparisons test **p<0.01 compared to WT). (B) *Gpr151*-KO mice earned significantly more nicotine infusions at the higher nicotine dose (0.4 mg/kg) (n=4-8, RM 2-way ANOVA, Bonferroni’s multiple comparisons test **p<0.01 compared to WT). (C) Total amount of nicotine self-administered at each dose (n=4-8, RM 2-way ANOVA, Bonferroni’s multiple comparisons test ***p<0.001 compared to WT). *Gpr151*-KO mice self-administer a significantly higher amount of nicotine at the highest 0.4 mg/kg/session dose. (D) Re-expression of GPR151 in the Mhb and IPN terminals of *Gpr151*-KO mice injected with AAV2.1-*Gpr151* visualized by GPR151 immunoreactivity and comparison with *Gpr151*-KO mice injected with AAV2/1-EGFP control virus. Scale bar: 500 um. (E) At the highest nicotine dose (0.4mg/kg/session), *Gpr151*-KO mice injected with the AAV2.1-*Gpr151* rescue virus significantly decrease the amount of self-administered nicotine to similar levels as WT (left panel) (n=6-12, unpaired t-test **p<0.01). Data are represented as mean ± SEM. See Table S3 for details of statistical analysis.

Using the same paradigm but pairing the lever pressing to intravenous infusion of nicotine, we next investigated the role of GPR151 in nicotine reinforcement. As expected, WT mice responded for self-administered nicotine infusions according to a known inverted U-shaped dose–response curve (Fowler et al., 2011) (Figure 3B). Notably, *Gpr151*-KO mice self-administered far more nicotine at a high nicotine dose than WT littermate mice (Figure 3B,C,E). This pattern of responding for nicotine, particularly at higher unit doses of the drug, is similar to that previously reported in α5 nAChR subunit KO mice and rats in which α5 subunits are selectively knocked down in Hb-IPN (Fowler et al., 2011), suggesting reduced sensitivity to the aversive effects of high doses of nicotine. Re-expression of GPR151 in the MHb of *Gpr151*-KO mice reduced self-administration of the 0.4mg/kg nicotine dose to levels similar to WT mice (Figure 3D,E). Consistent with the locomotor behavioral tests (Figure 2A,B,C), both the reduction in the anorectic effects of nicotine and the fact that *Gpr151*-KO mice self-administer more nicotine at a high doses suggests that the KO mice are resistant to the aversive and malaise-inducing effects of nicotine. Altogether, these results demonstrate that GPR151 is key in the inhibitory control exerted by the MHb in limiting drug and food intake.

### GPR151 is localized to axonal and presynaptic membranes and synaptic vesicles of habenular terminals

To understand the mechanism by which GPR151 modulates the sensitivity to nicotine, morphine and cannabinoids that act on their cognate habenular receptors, we next determined its precise subcellular localization. To this end, we employed immuno-electron microscopy (iEM) to visualize axonal projections and synaptic terminals in the rostral portion of the IPN (Figure 4A), which is most intensely labeled by GPR151 (Broms et al., 2015) and contains the highest concentration of cholinergic and glutamatergic axonal terminals arising from habenular neurons (Frahm et al., 2015). GPR151 immunoreactivity was observed in axon terminals of WT mice (Figure 4B and Figure S8A-B), but not in *Gpr151*-KO mice (Figure 4C and Figure S8C-D), confirming the specificity of the antibody. As expected from the characteristic architecture of the IPN, quantitative analysis of terminals containing an active zone (AZ) showed that most synaptic contacts labeled by GPR151 were axo-dendritic (50 out of 53), 2 were axo-axonic and 1 was axo-somatic (Figure S8E). To exclude the possibility that loss of GPR151 could have altered the morphology of IPN synapses we performed a comparative analysis of different synaptic markers in WT and *Gpr151*-KO terminals. No significant differences were observed in the area of the synaptic terminal, length of the AZ, diameter and density of SVs, and distance of SVs to the AZ (Figure 4D-H).

**Figure 4.**
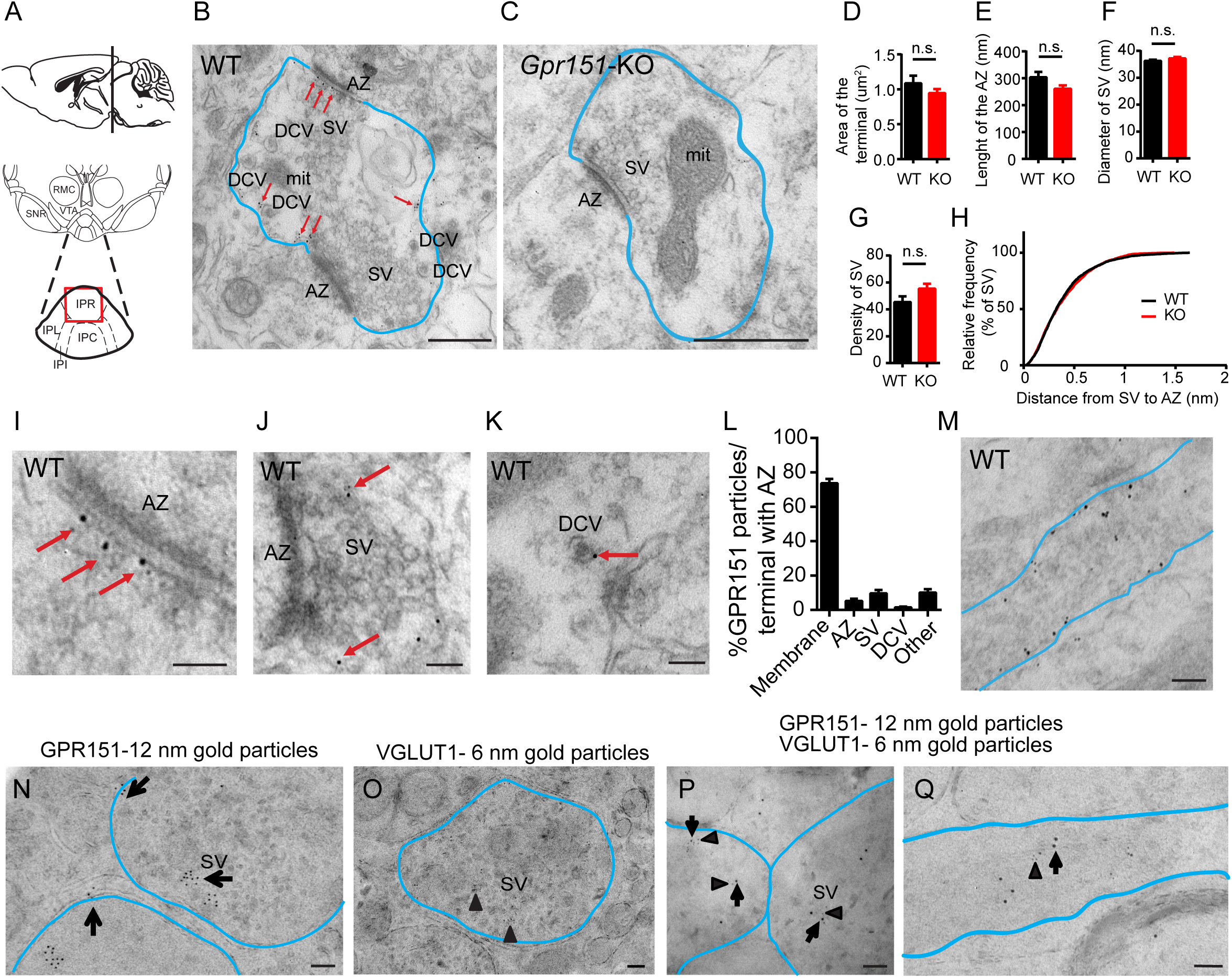
GPR151 is localized at MHb synaptic inputs to IPN. (A) Schematic of the brain area dissected for EM. (IPR: rostral, IPL: lateral, IPI: intermediate, IPC: central subnucleus of the IPN). (B) Representative micrograph showing GPR151 immunogold particles at the presynaptic terminals of WT mice. The presynaptic terminal is delineated in blue. Red arrows point to GPR151 nanogold particles located over the active zone (AZ) or at the membrane. In addition to synaptic vesicles (SV), the terminals contained dense core vesicles (DCV). Scale bar: 500 nm. (mit: mitochondria). (C) Absence of GPR151 immunogold particles in *Gpr151*-KO mice. Scale bar: 500 nm. (D-H) WT and *Gpr151*-KO mice show no significant differences in synaptic terminal area, length of the active zone and synaptic vesicle diameter, density, and distance to the AZ (n≥45, unpaired t-test, Table S3). (I-K) Representative micrographs showing GPR151 immunogold particles at the AZ (I), SVs (J) and DCV (K). Scale bar: 100 nm. (L) Quantitative analysis shows that within a synaptic terminal, GPR151 is mostly at the membrane (73.63% ± 2.54) but can also be found at the AZ (5.19 % ± 1.33) and, in association with SV (9.66% ± 1.890); DCV is (1.39% ± 0.51); and other structures (10.11% ± 1.95) (n=50 terminals, total of 999 particles). Data are represented as mean ± SEM. (M) Transversal section of an habenular axon showing GPR151 immunogold particles at the plasma membrane and along microtubules. Scale bar: 100 nm. (N-O) Immunogold particles for GPR151 (12 nm, arrows) and VGLUT1 (6 nm, arrowheads) are localized at presynaptic membranes and SVs in single (N,O) and double (P,Q) post-embedding EM experiments. Scale bar: 100 nm. Data are represented as mean ± SEM. See Table S3 for details of statistical analysis.

The majority of GPR151 nanogold particles (73.63%) were located at the presynaptic plasma membrane (Figure 4B and 4L), at the active zone (AZ) (5.19%; Figure 4B, 4I and 4L) and in association with synaptic vesicles (SV) (9.66% Figure 4J and 4L) and dense core vesicles (DCV) (1.39%, Figure 4K-L). SVs and DCVs labeled with GPR151 were mainly localized further than 200 nm from the AZ (Figure S8F-G). The distribution of GPR151 was similar in presynaptic terminals without an AZ in the plane of the picture (Figure S8B and S8H). GPR151 nanogold particles were also found along the habenular axonal projections, mostly at the membrane but also along the microtubules inside the axons (Figure 4M and Figure S9A-B). This is interesting in light of recent findings that nAChRs are also expressed along Hb axons in the fasciculus retroflexus, where they can be activated by nicotine (Passlick et al., 2018).

Next, we confirmed that GPR151 is at SVs by double postembedding iEM staining with the vesicular glutamate transporter 1 (VGLUT1) (Figure 4N-Q, Figure S9C-E). Out of 70 synaptic terminals analyzed, 87% (n=61) were labeled with both VGLUT1 and GPR151, while just 10% (n=7) were labeled with GPR151 only and 3% (n=2) were labeled with VGLUT1 only (Figure S9F). This indicates that around 90% of GPR151 habenular terminals at the IPR are glutamatergic. GPR151 and VGLUT1 were found very close to each other in association with synaptic vesicles (Figure 4P) and along the microtubules within the axons being transported towards the terminals (Figure 4Q and S9G-H). In summary these experiments identify GPR151 as a novel and specific component of presynaptic habenular terminals in the IPN. The fact that *Gpr151*-KO mice have presynaptic terminals with normal morphology and SVs distribution argues against a role for GPR151 in synaptic scaffolding or cytoskeleton organization or in brain development. However, GPR151 localization at the synaptic and perisynaptic membrane and SVs suggests that GPR151 may modulate synaptic activity.

### GPR151 contributes to synaptic fidelity, strength and plasticity

To explore the possibility that GPR151 modulates the synaptic activity of habenular terminals, we tested whether optogenetic activation of MHb terminals could reveal differences between WT and *Gpr151*-KO mice. Since the majority of GPR151 expressing neurons (70%) are cholinergic MHb neurons (Figure S10), we crossed *Gpr151*-KO mice with transgenic mice expressing the light-gated cation channel channelrhodopsin-2 (ChR2) in cholinergic neurons (*Chat*-ChR2 mice) (Ren et al., 2011) and recorded IPN neurons from offspring mice (Figure 5A). eEPSCs were evoked by trains of 5 ms blue light pulses (Figure 5B) and the fidelity of transmission was calculated as the percentage of light pulses that resulted in detectable postsynaptic currents per second during 1 min (Figure 5C). We observed increased fidelity rates in MHb-IPN synapses of *Chat*-ChR2x*Gpr151*-KO mice compared to *Chat*-ChR2 mice (Figure 5C) and compared to *Chat*-ChR2x*Gpr151*-KO mice injected with the rescue virus (AAV2/1-*Gpr151*) specifically in MHb (Figure 5D). However, the amplitude of the light evoked EPSCs was smaller in *Chat*-ChR2x*Gpr151*-KO (Figure 5E). Next we used the Paired Pulse Ratio (PPR) to measure presynaptic release probability upon light stimulation. At basal conditions, the PPR was similar between genotypes (Figure 5F). However, nicotine application in *Chat*-ChR2 slices increases the probability of the initial presynaptic release of the readily releasable pool of vesicles and therefore reduces the PPR (Figure 5G, 5I). In contrast, the PPR of *Chat*-ChR2x*Gpr151*-KO does not change upon nicotine application (Figure 5H-I). This is consistent with the fact that GPR151 regulates behavioral actions to nicotine, The increased synaptic fidelity, smaller EPSC and unchanged PPR upon nicotine administration in *Chat*-ChR2x*Gpr151*-KO point to a critical role of GPR151 in maintaining the readily releasable pool (RRP) of synaptic vesicles in habenular terminals in the IPN.

**Figure 5.**
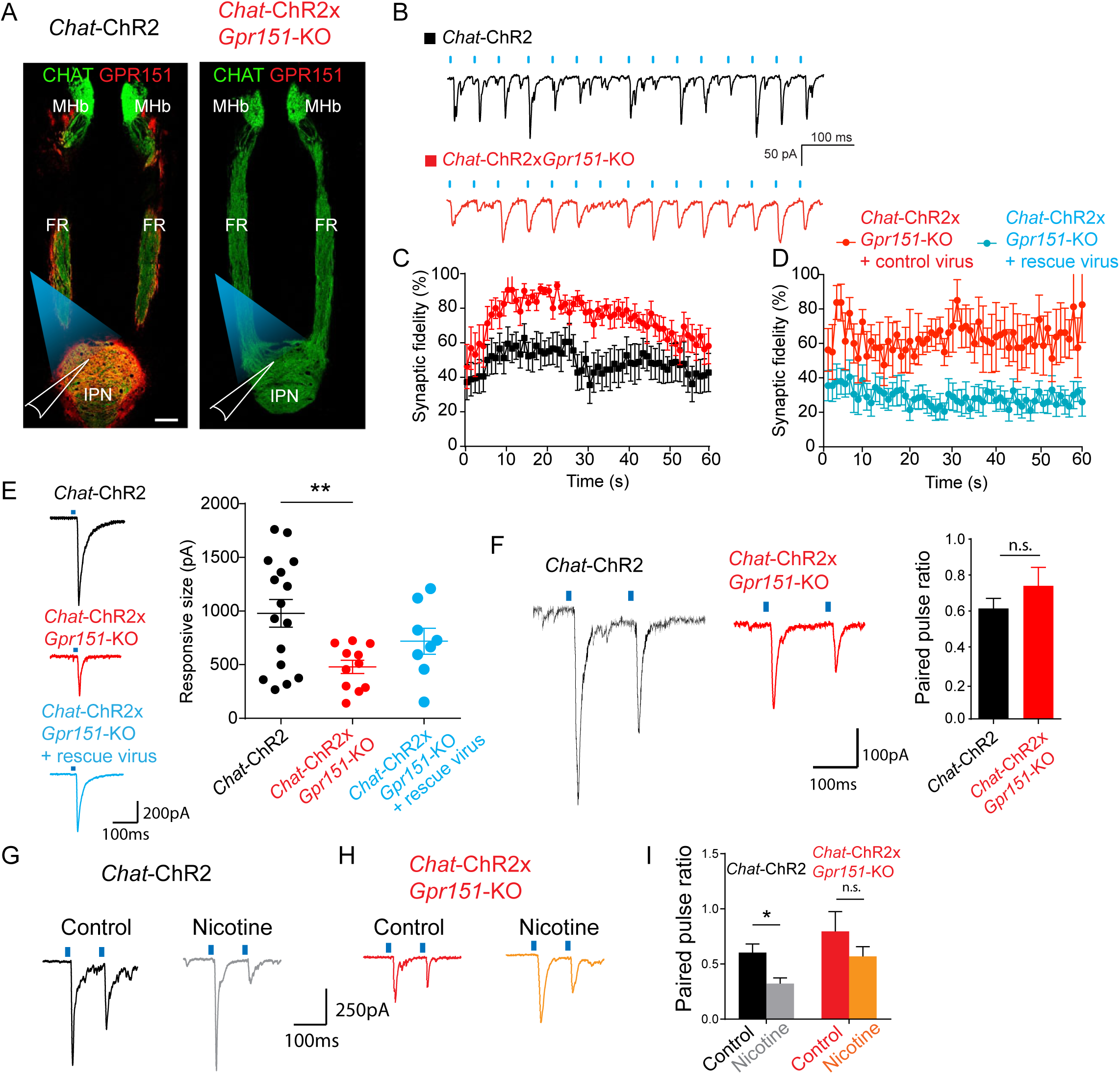
GPR151 contributes to synaptic plasticity. (A) *Chat*-ChR2-EYFP mice were crossed to *Gpr151*-KO for optogenetic recordings. Angled brain sections showing ChR2 (green) and GPR151 (red) in MHb-FR-IPN axis in *Chat*-ChR2 and loss of GPR151 signal in *Chat*-ChR2x*Gpr151*-KO. *Chat*-ChR2 terminals in the IPN were optogenetically stimulated and postsynaptic IPR neurons were recorded. Scale bar: 100 µm. (B) Voltage clamp recordings showing the fidelity of the synaptic responses to blue laser stimulation at 20Hz (5 ms pulse width). (C) Average synaptic fidelity was calculated as the percentage of light pulses that lead to successful transmissions for every second during 1 min at 20Hz stimulation. *Chat*-ChR2x*Gpr151*-KO mice (red) showed higher synaptic fidelity compared to *Chat*-ChR2 mice (black) (n=8-9, RM 2-way ANOVA, F_genotype_(1,15)=6.707, p=0.0205). (D) Viral re-expression of GPR151 in the MHb of *Chat*-ChR2x*Gpr151*-KO mice (blue) decreases synaptic fidelity compared to *Chat*-ChR2x*Gpr151*-KO mice injected with control virus (red) (n=4-10, RM 2-way ANOVA, F_genotype_(1, 12) = 8.379, p = 0.0135). (E) Synaptic strength measured as the amplitude of the first blue light evoked EPSC is reduced in *Chat*-ChR2x*Gpr151*-KO mice (red) compared to *Chat*-ChR2 mice (black) and partially restored by viral re-expression of GPR151 (blue) (n=8-16, one-way ANOVA, Tukey’s multiple comparisons test **p=0.0096 WT vs KO). (F) Example traces of paired pulse ratio (PPR) recordings in *Chat*-ChR2 (black) *Chat*-ChR2x*Gpr151*-KO (red) mice. The amplitude of the optically evoked EPSCs is reduced in KO as shown in (E) but the paired pulse ratio (PPR) was similar between genotypes (n=17-16, unpaired t-test, p=0.28). (G-H) Example traces of PPR after vehicle or nicotine (I) Nicotine-induced decreases in PPR were absent in *Chat*-ChR2x*Gpr151*-KO mice (n=17-8, Kruskall Wallis test, Dunn’s multiple comparisons test *p<0.05) Data are represented as mean ± SEM. See Table S3 for details of statistical analysis.

### GPR151 couples to the G-alpha inhibitory protein Gαo1 and associates with presynaptic components

To understand the signaling mechanisms through which GPR151 controls habenular neurons we sought to identify the G-alpha protein subunit groups (Gαs, Gαi, Gαq/11, or Gα12/13,) it uses for signal transduction. To accomplish this, we collected brain samples from the IPN of WT and *Gpr151*-KO mice, immunoprecipitated (IP) the protein extracts with GPR151 antibodies and performed mass spectrometry analyses of IP samples (Figure 6A, Figure S11A and Table S4). As shown in the volcano plot (Figure 6B), GPR151 is the most enriched protein in the immunoprecipitated fraction in WT mice, and it is not present in *Gpr151*-KO extracts, confirming the efficacy and specificity of the IPs. In addition, we identified 17 proteins that co-immunoprecipitated with GPR151 (Table S4). Among these, we were especially interested in the interaction with the G-alpha inhibitory subunit Gnao1 (protein name Gαo1), a heterotrimeric guanine nucleotide-binding G-proteins involved in intracellular signal transduction (Figure 6B). To confirm this interaction, we performed immunoprecipitations and Western blot analysis of IPN extracts from WT and KO mice. As shown in Figure 6C and 6D, GPR151 associates with Gαo1 but not with the G-alpha stimulatory subunit (Gαs). Consistently, the TRAP data shows that Gαo1 is the most abundant of the inhibitory Gi/o proteins in cholinergic MHb neurons (Table S2). These results indicate that GPR151 couples to the G-alpha inhibitory pathway, which is known to inhibit adenylyl cyclase activity and therefore decrease intracellular cAMP levels.

**Figure 6.**
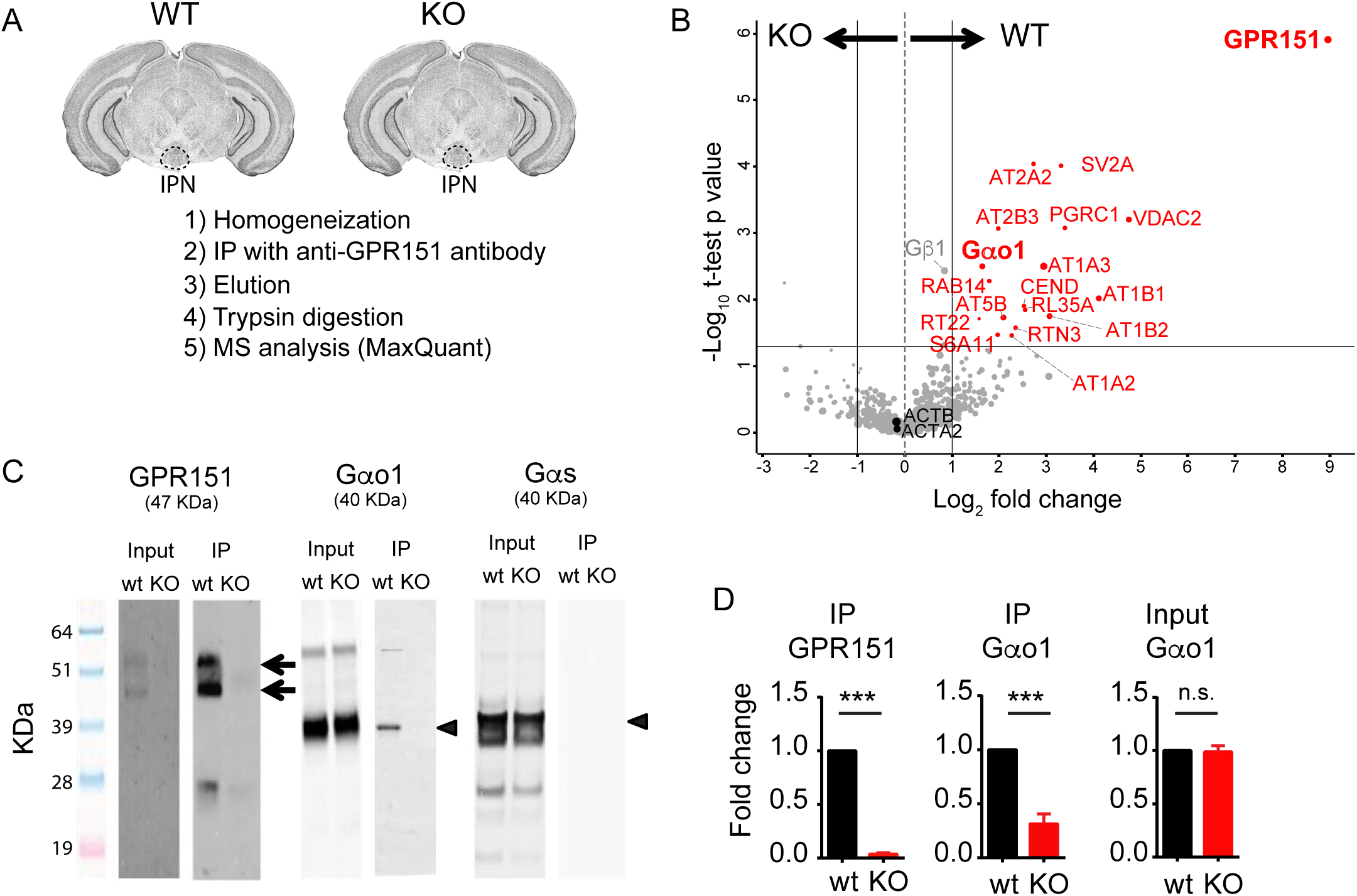
GPR151 couples to the G-alpha inhibitory subunit Gα_o1_ and co-immunoprecipitates with pre-synaptic regulators. (A) Outline of the co-immunoprecipitation (co-IP) and Mass Spectrometry (MS) experiments (n=5 biological replicates for WT and n=5 for KO. For each replicate, 15 IPN were dissected). (B) Volcano plot of proteins that co-IP with GPR151 from the IPN of WT mice (right) and *Gpr151*-KO (left) IPN samples. Log2 ratios are plotted against the adjusted negative log10 P values. Significant different proteins (p<0.05) found in WT samples are labeled in red. The size of the circles indicates an estimate of protein amount. P values were obtained using Student t-test. As expected GPR151 was the most enriched protein in the IP fraction of WT mice and was not present in KO mice. Other co-IP proteins include Gα_o1_, ATPases and SV2A. Control proteins not enriched in WT (log2<1) include Gβ_1_, ACTB1 and ACTA2 indicated in grey. (C-D) Western blot confirmation that GPR151 IPs with GPR151 in WT mice (double arrow) and that Gα_o1_ (arrowhead) but not Gα_s_ IPs with GPR151 in WT mice (IP GPR151 WT: 1 ± 0, KO: 0.03 ± 0.01, n=5, unpaired t-test p < 0.0001; IP Gαo1/2 WT: 1 ± 0, KO: 0.31 ±0.09, n=5, unpaired t-test p < 0.0001; Input Gαo1/2 WT: 1 ± 0, KO: 0.98 ± 0.05, n=5, unpaired t-test p=0.84). Data are represented as mean ± SEM.

Other proteins co-immunoprecipitated with GPR151 include synaptic vesicle proteins (Synaptic vesicle glycoprotein 2A (SV2A) and Voltage-Dependent Anion Channels (VDAC2)) (Morciano et al., 2009; Takamori et al., 2006) and proteins potentially involved in regulating the functional dynamics of the pre-synapse (including the sodium/potassium ATPase composed of AT1A2 and AT1A3 (catalytic α-subunits), AT1B1 and AT1B2 (structural β-subunits)) the calcium ATPase (AT2A2 and AT2B3), and the H+ transporting ATPase (AT5B) (Taruno et al., 2012; Zanni et al., 2012); (Figure 6B and Table S5, Figures S12, S13, S14 of GO analysis). Association of GPR151 with these proteins in mass spectrometry is consistent with immunoEM localization at the synaptic vesicle and with our electrophysiological findings demonstrating regulation of vesicular release.

### GPR151 regulates cAMP levels

Given its interaction with Gαo1, we next analyzed the possibility that GPR151 modulates cAMP levels. We conducted cAMP assays of IPN samples of WT and *Gpr151*-KO mice, as well as WT mice injected into the MHb with scramble shRNA or *Gpr151*-shRNA for downregulation (Figure 7A). Consistent with GPR151 coupling to Gαo1, we observed that cAMP levels are higher in *Gpr151*-KO IPN homogenates (Figure 7B). In addition, we observed a strong trend of increased cAMP levels in *Gpr151*-downregulated IPN (Figure 7B). Given that cAMP is an important regulator of neurotransmission of MHb cholinergic synapses (Hu et al., 2012), we performed electrophysiological recordings in *Chat*-ChR2 and *Chat*-ChR2x*Gpr151*-KO upon light stimulation as explained for Figure 5. As expected, addition of forskolin increased the amplitude of light evoked EPSCs in *Chat*-ChR2 mice (Figure 7C). However, the amplitude was not altered in *Chat*-ChR2x*Gpr151*-KO upon forskolin application and remained smaller than WT levels (Figure 7C). This suggests that the absence of GPR151 compromises the coupling of cAMP signaling to the neurotransmitter release machinery in habenula neurons.

**Figure 7.**
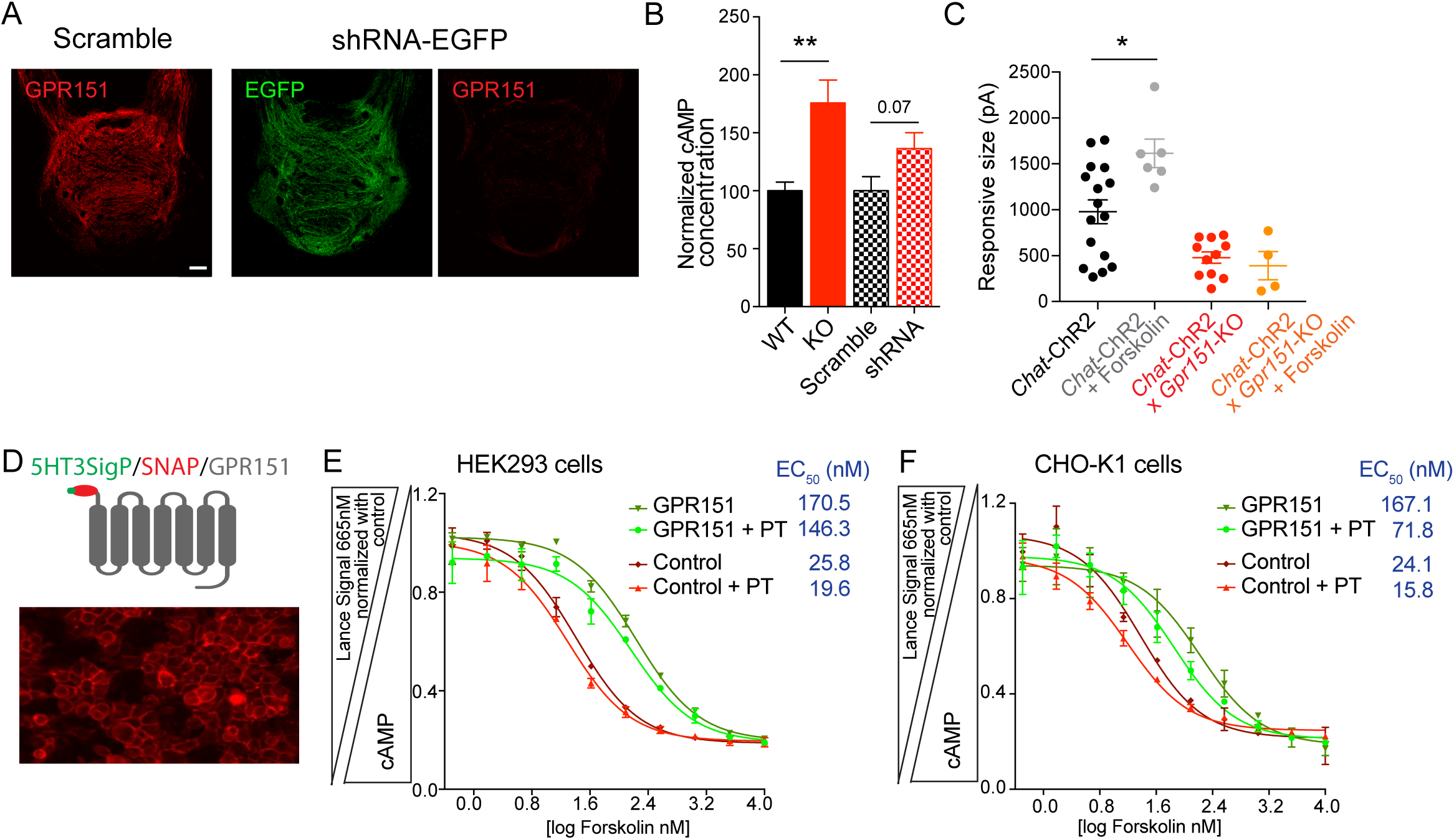
GPR151 regulates cAMP levels. (A) Knockdown of *Gpr151* in IPN after injection of LV-shRNA-GFP in the MHb of WT mice in comparison to mice injected with LV-scramble. Scale bar: 100um. (B) *Gpr151*-KO and shRNA-injected mice have higher cAMP levels in the IPN than WT and LV-scramble control mice (KO vs WT: n=15, unpaired t-test p=0.0016, shRNA3 vs scramble n=8-6, unpaired t-test p=0.07). (C) Forskolin increases the amplitude of light evoked EPSCs in *Chat*-ChR2 neurons (n=16-6, one-way ANOVA, Bonferronís multiple comparison test *p=0.0149), but has no effect in *Chat*-ChR2x*Gpr151*-KO neurons (n=11-4, one-way ANOVA, Bonferronís multiple comparison test p>0.99). (D) The signal peptide of the 5HT3 receptor and a SNAP-tag encoding region were used to express and monitor expression of human GPR151 in stable cell line. Staining with the cell impermeable SNAP dye 549 shows GPR151 expression at the surface in HEK293 stable cell clone. (E-F) LANCE cAMP competition assay in HEK293 (E) and CHO-K1 cells (F) under forskolin stimulation (log forskolin nM) shows higher cAMP levels in parental control cells (red) than in GPR151-expressing cells (green). Application of Pertussis toxin (PT) that inactivates G-alpha inhibitory subunits including Gα_o1_ (present in CHO-K1 but not in HEK293) decreases the EC50 in CHO-GPR151 but not in HEK293-GPR151 cells (n=2 technical triplicates for each response curve, one-way ANOVA for EC_50_ values, Bonferroni’s multiple-comparison test, **p=0.0085 HEK293-Control vs HEK293-GPR151, p>0.99 HEK293-Control vs HEK293-Control+PTX, p>0.99 HEK293-GPR151 vs HEK293-GPR151+PTX, ****p<0.0001 CHO-K1-Control vs CHO-K1-GPR151, p>0.99 CHO-K1-Control vs CHO-K1-Control+PTX, ***p=0.0004 CHO-K1-GPR151 vs CHO-K1-GPR151+PTX). Data are represented as mean ± SEM. See Table S3 for details of statistical analysis.

To further analyze the role of GPR151 in cAMP modulation and validate its interaction with Gαo1, we generated two stable cell lines expressing GPR151 with an N-terminus 5HT3 signal peptide to enhance membrane expression (Wellerdieck et al., 1997; Wetzel et al., 1999) plus a SNAP-tag to monitor surface expression (Ward et al., 2011) in HEK293 and CHO-K1 cells. CHO-K1 cells were used because they express high levels of Gαo1. In contrast, HEK293 cells do not express Gαo1, but express Gαz, which is a member of the same Gαi/o family (Baycin-Hizal et al., 2012; Doi et al., 2016; Thakur et al., 2011). GPR151 expression at the membrane was validated by SNAP and GPR151 immunostaining and western blot (Figure 7D and S15). Next we measured cAMP levels upon forskolin stimulation using the LANCE cAMP assay. The EC_50_ of forskolin of GPR151-expressing HEK293 (170.5nM) or CHO-K1 (167.1nM) cells were significantly higher than their parental cells, HEK293 (25.8nM) and CHO-K1 (24.1nM) (Figure 7E-F), suggesting that GPR151 couples to Gαi/o proteins and that its constitutive activity buffers the effect of forskolin.

Next we treated the cells with 250 ng/ml pertussis toxin (PT) overnight, which inactivates the G proteins by catalyzing the ADP-ribosylation of the αsubunits of the heterotrimeric Gαi/o protein family, with the exception the Gαz isoform (Fong et al., 1988; Matsuoka et al., 1988). PT had a strong effect on cAMP production stimulated by forskolin in GPR151-expressing CHO-K1 cells (Figure 7F). The EC_50_ of forskolin decreased from 167.1nM to 71.82nM indicating that PT blocks a large portion of the inhibitory response mediated by Gαo. The impact of PT in GPR15-expressing HEK293 cells is not as strong as in CHO-K1 cells (Figure 7E). The EC_50_ of forskolin in HEK293-GPR151 expressing cells decreased from 170.5 nM to 146.3 nM upon PT treatment, most likely because HEK293 cells have Gαz, which is insensitive to PT.

Together these results provide evidence that GPR151 couples to Gαo and decreases cAMP levels. In agreement with the lower cAMP levels observed in the IPN of WT mice compared to *Gpr151-*KO mice, GPR151 was observed to be constitutively active in both cell lines analyzed. This constitutive activity of GPR151 is actually not unexpected since it lacks the DRY motif which plays an essential role in class A-GPCR’s function (Wess, 1998). Mutations or deletion of the DRY motif lead to the formation of active GPCR conformations (Rosenkilde et al., 2005; Wess, 1998). Moreover, GPCRs without the DRY motif have been shown to be both constitutively active and ligand-inducible; and that inserting the DRY motif decreases its constitutive activity (Rosenkilde et al., 2005). Our data suggest that GPR151 could be also functioning via both a constitutive and a ligand-inducible mechanism.

## DISCUSSION

The conservation of the habenula across vertebrates and its emerging role in processing reward-related and aversive signals, including drug abuse responses, points to the need to further characterize this brain circuit and identify modulatory mechanisms that may provide common avenues for the development of new approaches towards the treatment of addiction and other disorders of the “reward” circuitry. Our previous studies showed the specific expression of the orphan receptor GPR151 in the habenula of rats, mice and zebrafish. Here, we establish conservation of the habenular-enrichment of this receptor in human brains, define its function in regulating habenular neurons, and reveal its critical role in regulating behavioral response to addictive drugs across different classes. More specifically, we demonstrate that GPR151 is present in presynaptic termini of rodent and human habenular neurons, and that it couples to Gαo to regulate cAMP levels, synaptic fidelity and neurotransmission. We also show that neurons in the medial habenula co-express GPR151 with nAChRs, MOR and CB1R, and that loss of GPR151 results in alterations in behaviors associated with nicotine, opioid and cannabinoid use. Thus, our data identify the MHb/IPN projection as a common pathway controlling addictive-relevant behaviors in response to major classes of drugs of abuse, and they demonstrate that this important circuit is modulated by the orphan receptor GPR151.

We found that GPR151 is selectively enriched in habenular presynaptic structures in the IPN. Several GPCRs, including dopamine D2, muscarinic M2 and M4, adrenergic, cannabinoid and opioid receptors have been localized at the presynapse where they inhibit evoked synaptic transmission (Betke et al., 2012). GPCRs directly at the active zone regulate fast transmission by local action of the dissociated G beta-gamma complex (Gβγ) on exocytosis, while GPCRs distant from the AZ modulate neurotransmission through second-messenger cascades (Betke et al., 2012). Following with this distinction, GPR151, which is mainly at the presynaptic plasma membrane (73% gold particles), most likely regulates synaptic transmission through cAMP second messenger cascades rather than through Gβγ. However, our mass spectrometry analysis revealed that GPR151 binds to Gαo1 and Gβ1 subunits, shown to localize to SVs and DCVs in neurons (Ahnert-Hilger et al., 1994; Takamori et al., 2006). The fact that GPR151 can be detected at the AZ, in SVs and in DCVs (17% of immunoparticles) and that it immunoprecipitates with Gαo1 and Gβ1, suggests the presence of preassembled GPR151/G-protein signaling complexes in vesicles that can be transported to the plasma membrane upon neuronal stimulation. Interestingly, GPR151 can be activated by acidic conditions in vitro (Mashiko et al., 2019) suggesting the possibility that GPR151 is activated by acidic SVs which release protons during neurotransmission (Traynelis and Chesler, 2001). Acidification also occurs after neuronal injury and acidosis induces pain and hyperalgesia (Ortega-Ramirez et al., 2017). An increase of *Gpr151* mRNA (but no GPR151 protein) has been detected after nerve ligation in dorsal root ganglia (Holmes et al., 2017) and spinal cord (Jiang et al., 2018). However, diminished allodynia in *Gpr151*-mutant mice after neuropathic pain was only found in one of the studies (Jiang et al., 2018), suggesting that further studies will be required to examine its role in pain.

Consistent with the EM analyses, GPR151 co-immunoprecipitates with proteins localized at presynaptic terminals and at SVs, again suggesting that GPR151 preassembles not only with G-alpha and beta proteins but also with SV components during its translocation to the plasma membrane. Importantly, GPR151 did not co-immunoprecipitate with CB1R and MOR, which are found presynaptically in habenular terminals, and we did not observe differences in CB1R and MOR protein expression in *Gpr151*-KO mice. These data exclude the possibility that the reduced sensitivity to opioids and cannabinoids in *Gpr151*-KO mice is caused by stoichiometric changes in the formation of heteromeric complexes of these receptors or their chaperoning to the membrane. This is in contrast to findings with antipsychotic drugs, which have been reported to bind glutamate mGluR2/ serotonin 2AR heterocomplexes to result in behavioral alterations in response to a variety of pharmacological compounds (Fribourg et al., 2011).

Our studies suggest that enhanced cAMP levels in habenular terminals of *Gpr151*-KO mice are responsible for the behavioral changes observed. cAMP reduces synaptic failure rates (Huang and Hsu, 2006) and facilitates neurotransmission by increasing release probability at central excitatory synapses (Sakaba and Neher, 2001). This has also been reported in habenular neurons where depletion of presynaptic cAMP levels suppresses neurotransmission (Hu et al., 2012). Although enhanced synaptic fidelity improves the precision of information transfer through brain networks (Owen et al., 2013), in some cases, increased synaptic fidelity might be deleterious, for instance, when it is a consequence of chronic drug administration. Upregulation of the cAMP signaling pathway is a typical molecular adaptation in several brain regions following chronic morphine treatment (Bie et al., 2005), and activation of CB1R also modifies synaptic efficacy (Katona and Freund, 2008).

MOR and CB1R, like GPR151, interact with the Gαi/o class of adenylate cyclase inhibitory Gα proteins (Howlett and Fleming, 1984; Uhl et al., 1994). From these observations, it is reasonable to hypothesize that maintaining increased levels of cAMP at habenular terminals of *Gpr151*-KO blunts the response of these neurons to drugs of abuse. Thus, the picture emerging from our studies is that GPR151 constitutive activity in normal WT synapses maintains a low level of cAMP at habenular synapses that allows dynamic responses to fluctuation by drugs of abuse acting on habenular nicotinic, opioid and cannabinoid receptors. In *Gpr151*-KO mice, cAMP levels are very high and neither forskolin nor drugs of abuse can elicit changes of neurotransmitter release due to a ceiling effect. Since insensitivity to these drugs in *Gpr151*-KO mice leads to higher self-administration, the findings here suggest that future studies aimed at the identification of an agonist of GPR151 (that will further decrease the levels of cAMP at this synapse) could be useful to treat addiction. No studies in central synapses have evaluated whether nicotine increases cAMP levels, but it has been shown that nicotine acts presynaptically to augment the frequency of neurotransmitter release (Frahm et al., 2015). This suggests that in addition to changes in cAMP, other molecular mechanisms may determine release probability in habenular synapses, for instance through ATPases. Since GPR151 interacts with the sodium/potassium pump ATP1A3/B1 which contributes to precise and reliable transmission at glutamatergic synapses (Taruno et al., 2012), it is possible that GPR151 reduces synaptic strength during high-frequency stimulation by impairing ATP1A3/B1 trafficking or activity by regulating its phosphorylation (Poulsen et al., 2010). Further studies on this particular aspect would be key to understand how GPCRs contribute to ATPase function in synaptic fidelity.

Addictive drugs, although chemically different from one another, share common cellular and anatomical pathways. Two lines of investigation have led to this conclusion: 1) most of them activate the mesocorticolimbic dopaminergic (DA) reward circuitry, 2) each of them mimics or enhances the actions of one or more neurotransmitters in the brain that are involved in the control of the brain reward circuit (Koob and Volkow, 2010). Connections between the lateral habenula and the DA system have been established (either direct LHb-VTA, or via the RMTg-VTA-SNc) (Proulx et al., 2014), and between the MHb and VTA DA system via the IPN (Ables et al., 2017; Zhao-Shea et al., 2015). These observations suggest that drugs of abuse act at the MHb-IPN and mesocorticolimbic reward circuit to process both reward-related and aversive signals. Also we cannot exclude that *Gpr151-KO* behavioral effects may also involve the LHb since a subpopulation of LHb muscarinic M2 neurons is GPR151 positive. However consistent with our self-administration experiments, loss of GPR151 in habenula did not alter the rewarding effects of nicotine but abolished the inhibitory aversive effects of higher nicotine doses on brain reward. A balance between reward and aversion is also evident by the fact that not only tolerance, sensitization and self-administration responses are changed in *Gpr151*-KO mice, but also food intake by activation of CB1R and nAChRs. Animal studies and human imaging analyses have revealed that many psychoactive drugs act on the mesocorticolimbic dopamine system, a circuit that appears to be common to the rewarding effects of some drugs of abuse, as well as other reinforcing natural behaviors such as eating, thirst, gambling and sexual drive (Lammel et al., 2014; Lammel et al., 2012; Ostroumov and Dani, 2018). Feeding responses to drugs of abuse are also mediated by the mesolimbic DA circuit (Land et al., 2014; Nunes et al., 2013; Wise and McDevitt, 2018)and by hypothalamic POMC neurons (Koch et al., 2015; Mineur et al., 2011). Since neuroanatomical reciprocal connections exist between the IPN and hypothalamus (Groenewegen et al., 1986), it is possible that GPR151-mediated alterations of MHb-IPN neurotransmission impact feeding behavior by modulating postsynaptic responses of POMC neurons. It has been proposed that obesity and drug addiction arise from similar neuroadaptive responses (DiLeone et al., 2012; Kenny, 2011). Interestingly, UK biobank studies have recently associated GPR151 loss of function alleles to lower body mass index, warranting further studies on the role of *Gpr151* in feeding. However we only detected differences on feeding behavior when mice are treated with the cannabinoid agonist ACEA (not with saline control animals) or with nicotine. Our results highlight the habenula as a common key brain area in the development of addiction to some drugs of abuse and provide evidence of a role of this ancient epithalamic nucleus in feeding regulation and weight control. Thus our findings raise the possibility that malfunction of the habenula may predispose individuals to drug consumption and consequent changes in food intake that may in turn reinforce the use of drugs.

Finally, our studies suggest GPR151 as a candidate target for therapies preventing drug abuse because: 1) it is exclusively localized in axonal projections of habenular neurons, decreasing therefore the likelihood of developing side effects due to alterations in other brain circuits or in the peripheral nervous system, 2) it modulates neurotransmission and by that affects sensitivity to nicotine, morphine and cannabinoid-induced behaviors, 3) it belongs to the highly druggable Class A family of GPCRs and can be screened with cAMP assays, and 4) its pattern of expression is conserved in the human habenula-IPN axis. We hope our work stimulates interest in this orphan GPCR since the identification of a GPR151 agonist would be a very valuable therapeutic target to reduce the enormous adverse impact of drug abuse on public health.

## EXPERIMENTAL PROCEDURES

### Animals

All procedures involving mice were approved by the Rockefeller University and Mount Sinai Institutional Animal Care and Use Committee and were in accordance with the National Institutes of Health guidelines. See Supplemental Experimental Procedures for further details.

### Co-immunoprecipitations, western blot and proteomic analysis

Co-immunoprecipitations for mass spectrometry analysis were performed using M-270 Epoxy Dynabeads (Invitrogen) with three different GPR151 antibodies (45 µg aprox) coupled to 15 mg of beads per sample during 24 hr at 37 °C in 1.5 ml volume. Co-immunoprecipitations for western blot analysis were done using 50 µl of DynaBeads Protein G (Invitrogen) per sample and 6 µg anti-GPR151 antibodies. See Supplemental Experimental Procedures for further details and proteomic analysis.

### Immunohistochemistry of human and mouse brain samples and quantification analysis

Human brain tissues were obtained from the NICHD Brain and Tissue Bank from 5 donors, ranging between 22–52 years of age. Immunohistochemistry was performed in adult mice (8-12 weeks) as described in (Broms et al., 2015). See Supplemental Experimental Procedures for further details.

### Electron microscopy

Pre-embedding and post-embedding nanogold labeling was performed as in (Frahm et al., 2015). See Supplemental Experimental Procedures for details.

### TRAP and RNA-SEQ

Three biological replicates were used for TRAP analysis as described in (Mellen et al., 2012). Each replicate contained the habenulae from 5 CHAT-EGFP-L10a transgenic mice (males and females 8-12 weeks old). See Supplemental Experimental Procedures for details.

### Electrophysiological recordings

IPN neurons were patch-clamped at -70 mV. Presynaptic MHb fibers were excited for one continuous minute with a 473 nm blue light laser stimulation at 20 Hz, and a pulse length of 5 ms. The fidelity of the MHb-IPN transmission was calculated for every second as the number of successful transmissions, divided by 20 in percent. Kinetics of successful transmissions were calculated using MiniAnalysis (Synaptosot). See Supplemental Experimental Procedures for details.

### Behavioral analysis

All behavioral studies were conducted blind to the genotype of the tested mice and only male mice 8-16 weeks old were used. See Supplemental Experimental Procedures for details.

### Intravenous nicotine self-administration procedure

Mice were mildly food restricted to 85-90% of their free-feeding body weight and trained to press one of two levers in an operant chamber (Med-Associates Inc., St. Albans, VT) for food pellets under a FR5TO20 reinforcement schedule. When mice demonstrated stable responding (>25 pellets per session), they underwent jugular catheter implantation. Once stable responding was re-established after surgery, subjects were permitted to respond for intravenous nicotine infusions during 1 h daily sessions, 7 days per week, 3-5 days for each dose of nicotine in ascending order with saline last. In between each dose, subjects were permitted access to the training dose for at least 2 days or until their intake returned to baseline levels before being tested on the next dose. See Supplemental Experimental Procedures for more details.

### Statistical Analysis

Statistical analyses were performed with GraphPad Prism 6.0. Unpaired two-tailed Student t-tests, 2way ANOVA or Repeated Measures (RM) 2way ANOVA were used for analyzing most of the data as indicated in figure legends. See Table S3 for details of statistical analysis. Results are presented as means ± S.E.M.

## AUTHOR CONTRIBUTIONS

Study concept and design: PJK and II-T Acquisition of data: BA-F, AG, JLA, MD. Analysis and interpretation of data: BA-F, AG, JLA, PJK, and II-T. Writing of the manuscript: BA-F, PJK and II-T.

## ACKNOWLEDGMENTS

We thank Kunihiro Uryu and Nadine Soplop (Electron Microscopy Resource Center, Rockefeller University) for EM expertise. We thank Milica Tešić Mark and Henrik Molina (Proteomics Resource Center, The Rockefeller University) for mass spectrometry analysis, which acknowledges funding from the Leona M. and Harry B. Helmsley Charitable Trust. Human tissue was obtained from the NICHD Brain and Tissue Bank for Developmental Disorders at the University of Maryland, Baltimore, MD. We thank Awni Mousa for analysis of RNA-Seq data, Sylvia M. Lipford, Cuidong Wang and Juncheng Li for technical assistance and Kun Li and Nathaniel Heintz for critical discussions. A.G received a DFG fellowship (GO 2334/1-1). This work was supported by the Leona M. and Harry B. Helmsley Charitable Trust and Sohn Conferences Foundation for mass spectrometer instrumentation (The Rockefeller University Proteomics Resource Center), the Leon Black Family Foundation (II-T), NIDA (1P30 DA035756-01) (II-T), DA020686 (PJK)), and NIDA UG3 DA048385(PJK and IIT).

