## Supplementary figures and images for "The Habenular Receptor GPR151 Regulates Addiction Vulnerability Across Drug Classes"

### supplemental Fig 1

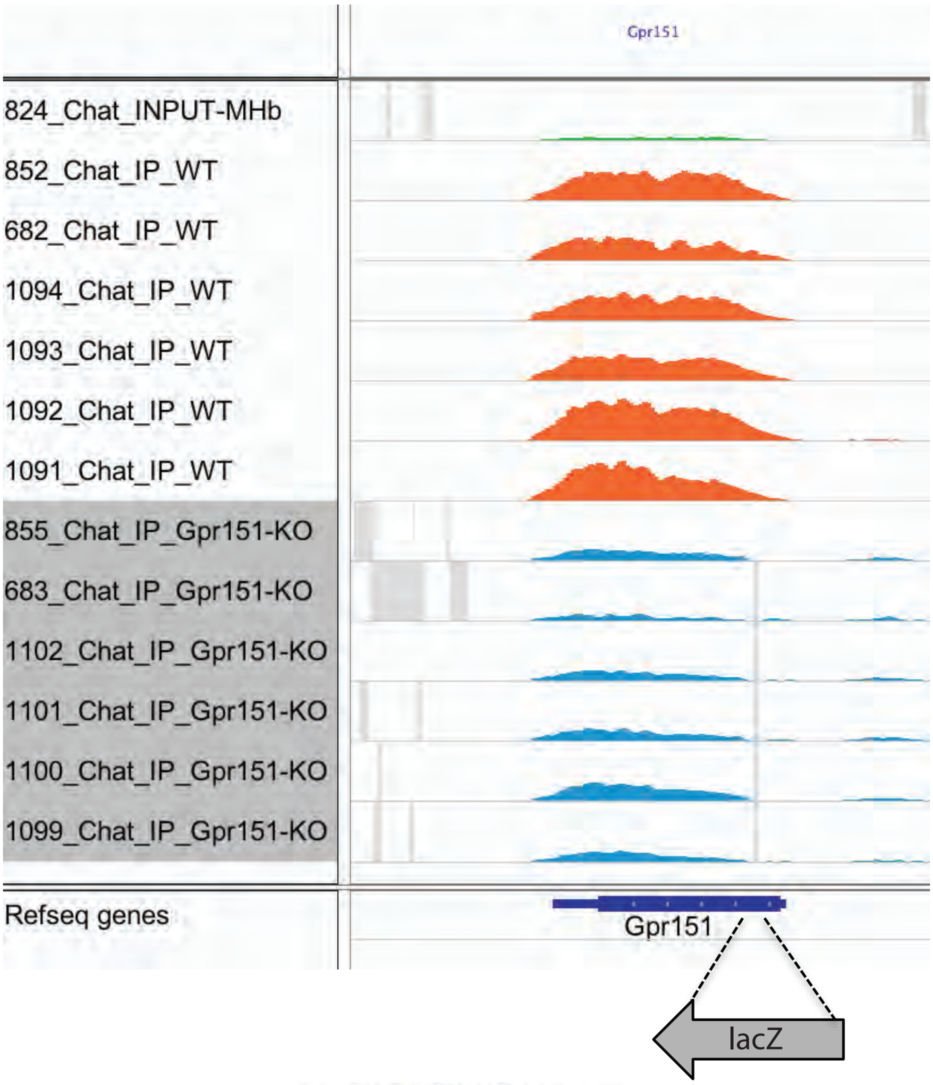

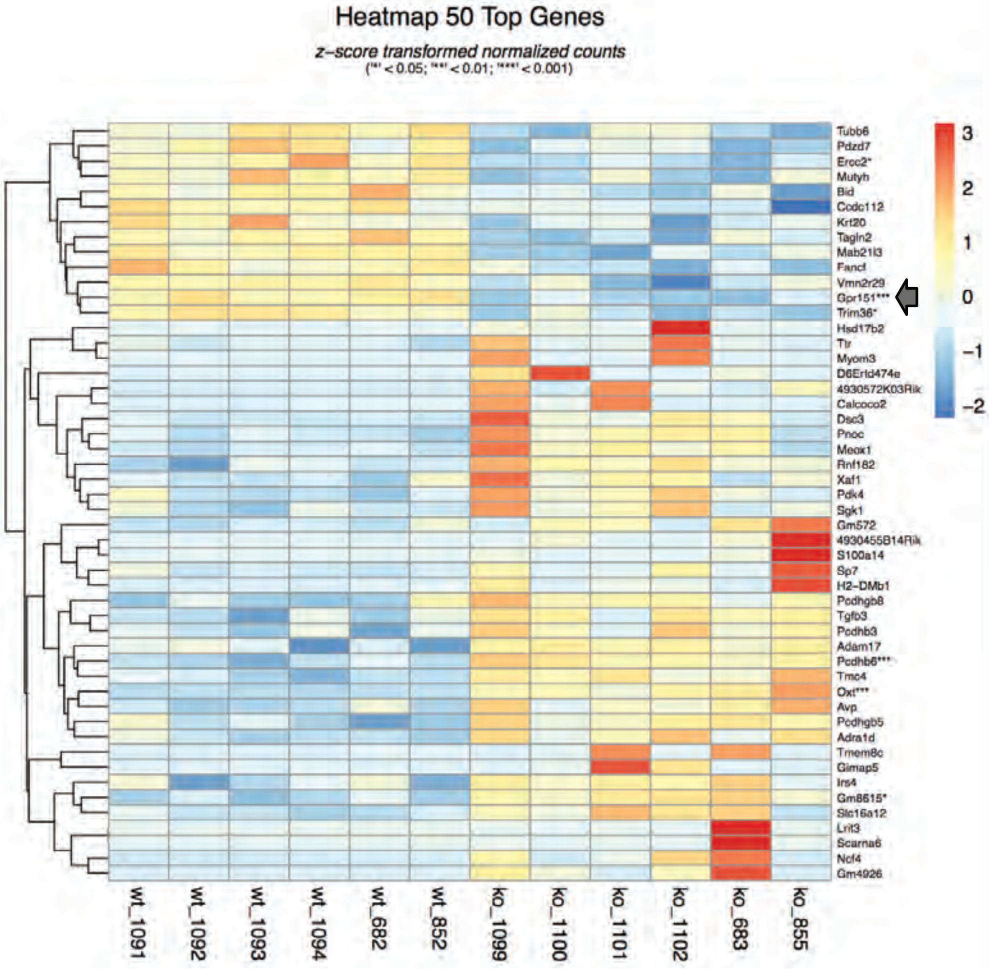

### supplemental Fig 2

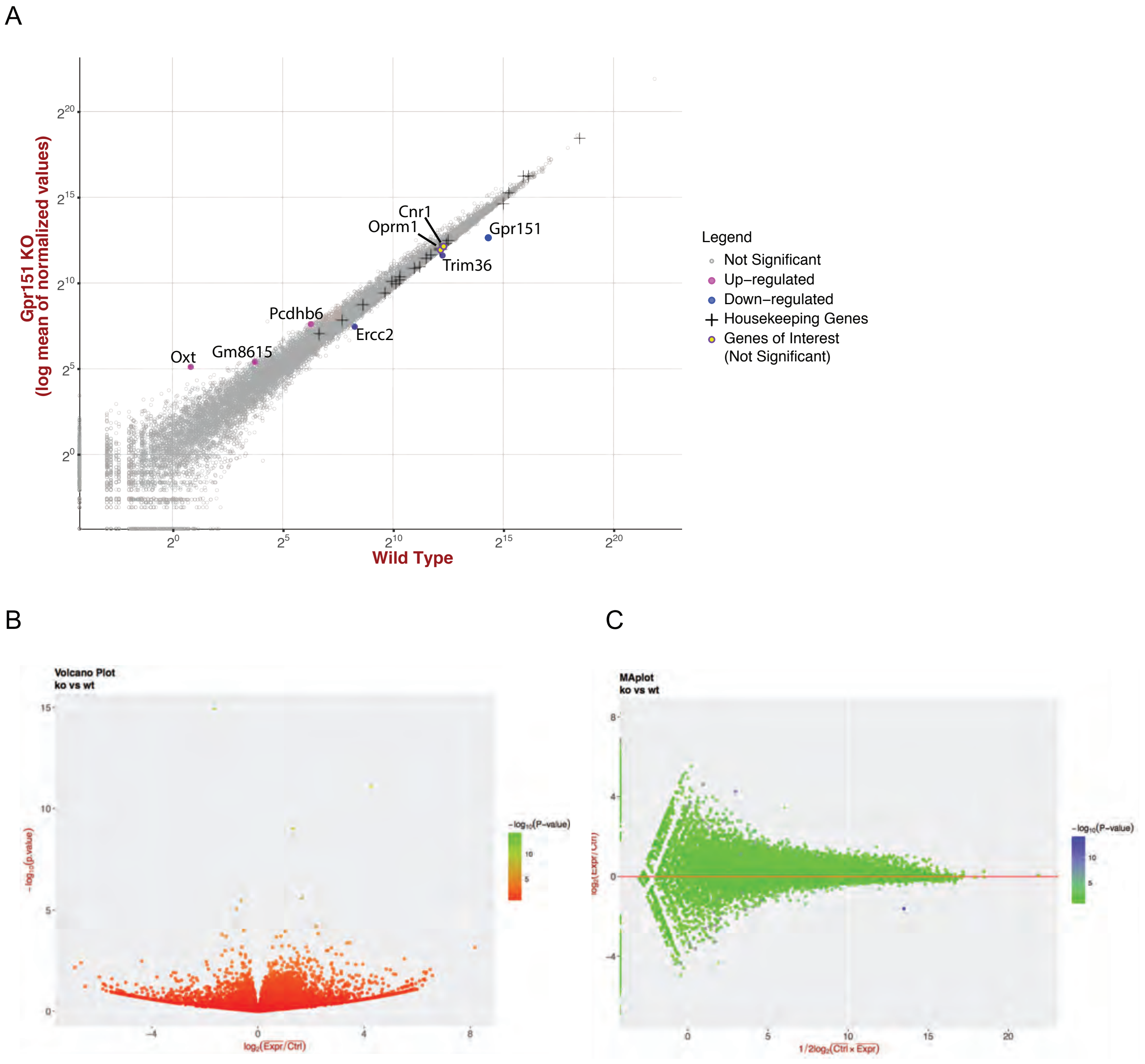

### supplemental Fig 3

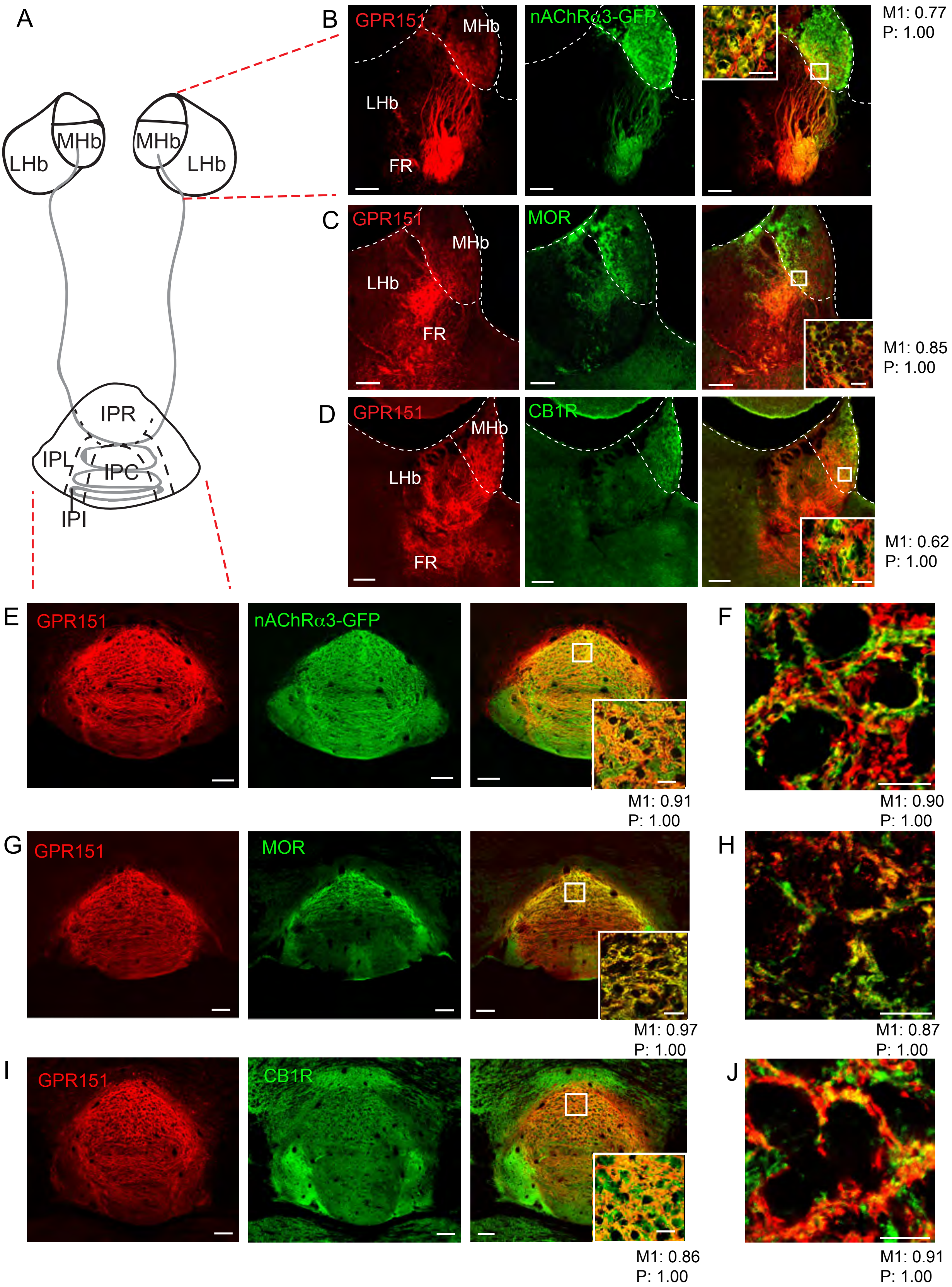

### supplemental Fig 4

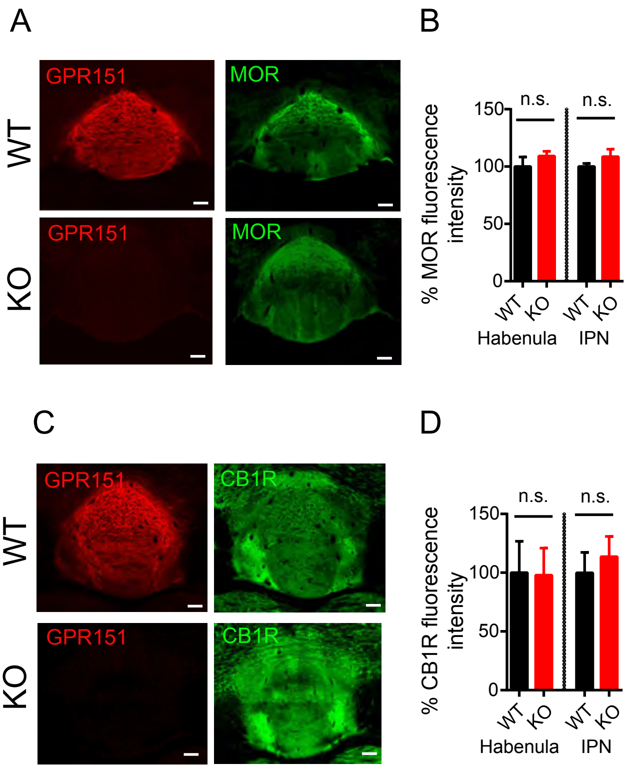

### supplemental Fig 5

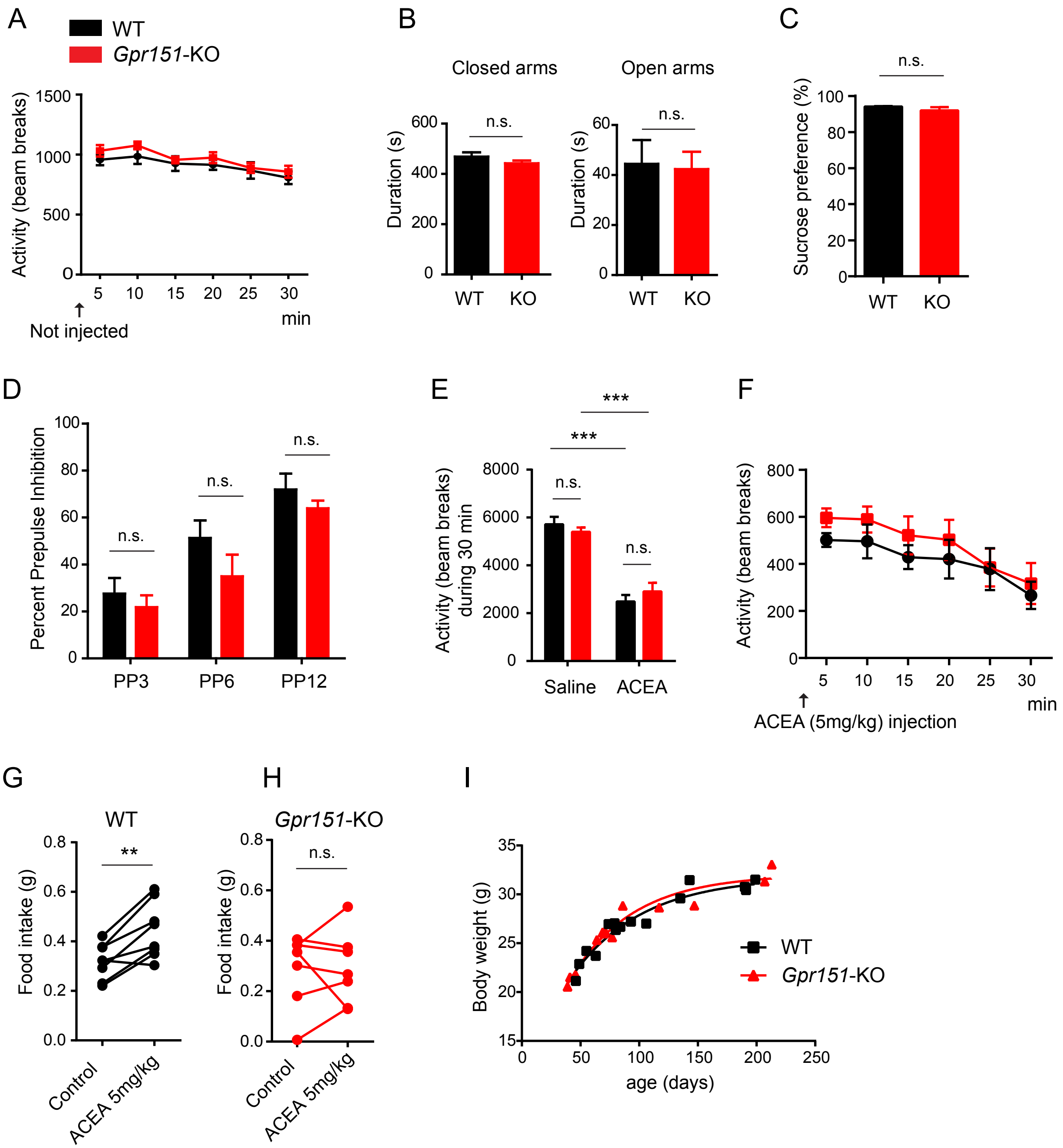

### supplemental Fig 6

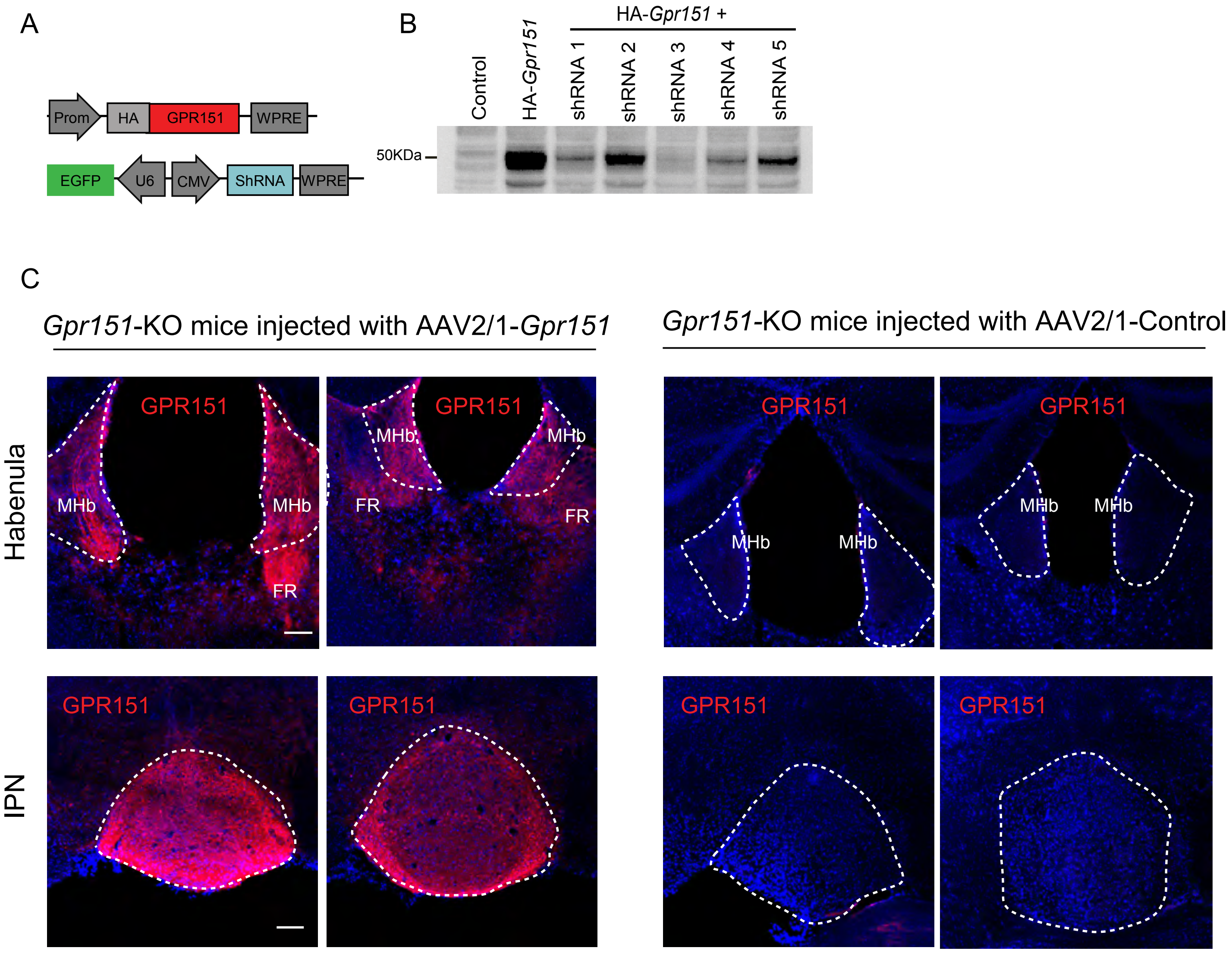

### supplemental Fig 7

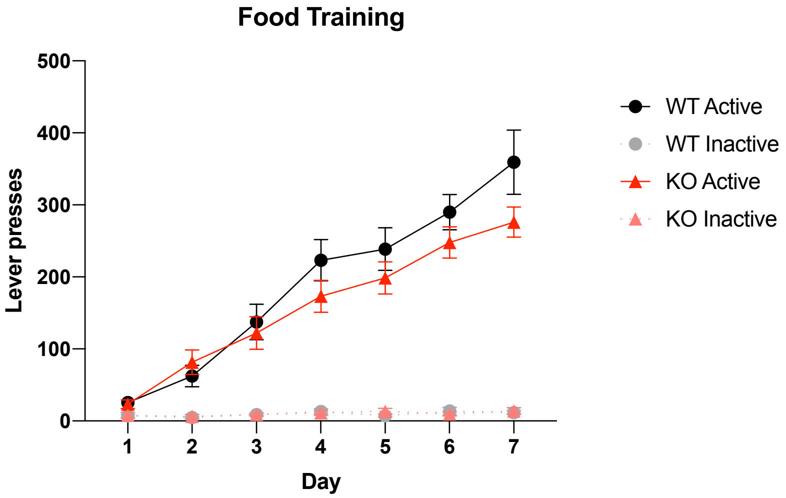

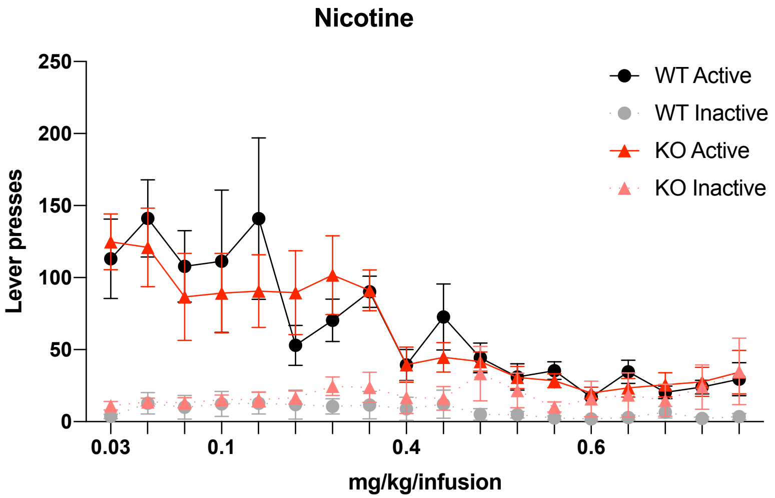

### supplemental Fig 8

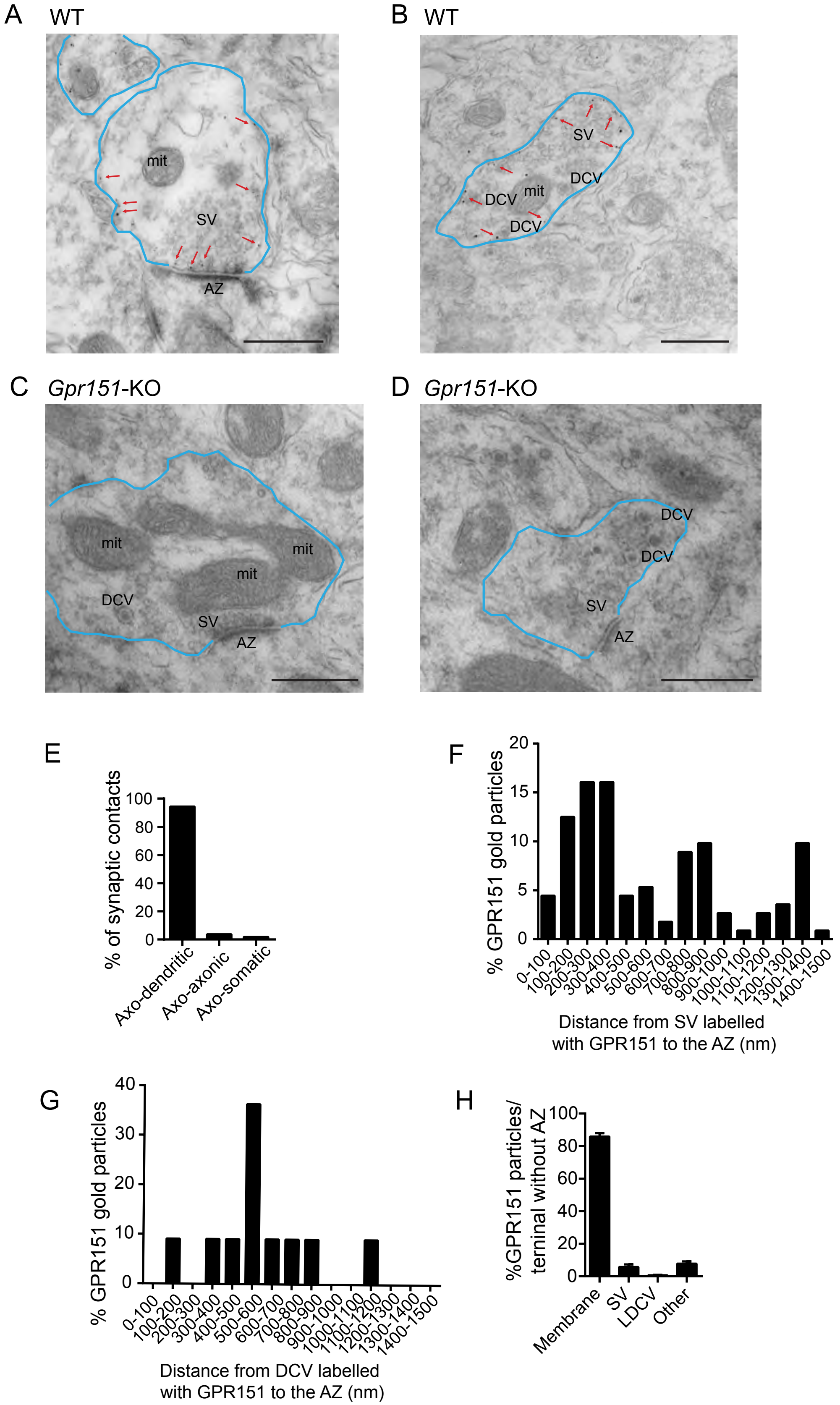

### supplemental Fig 9

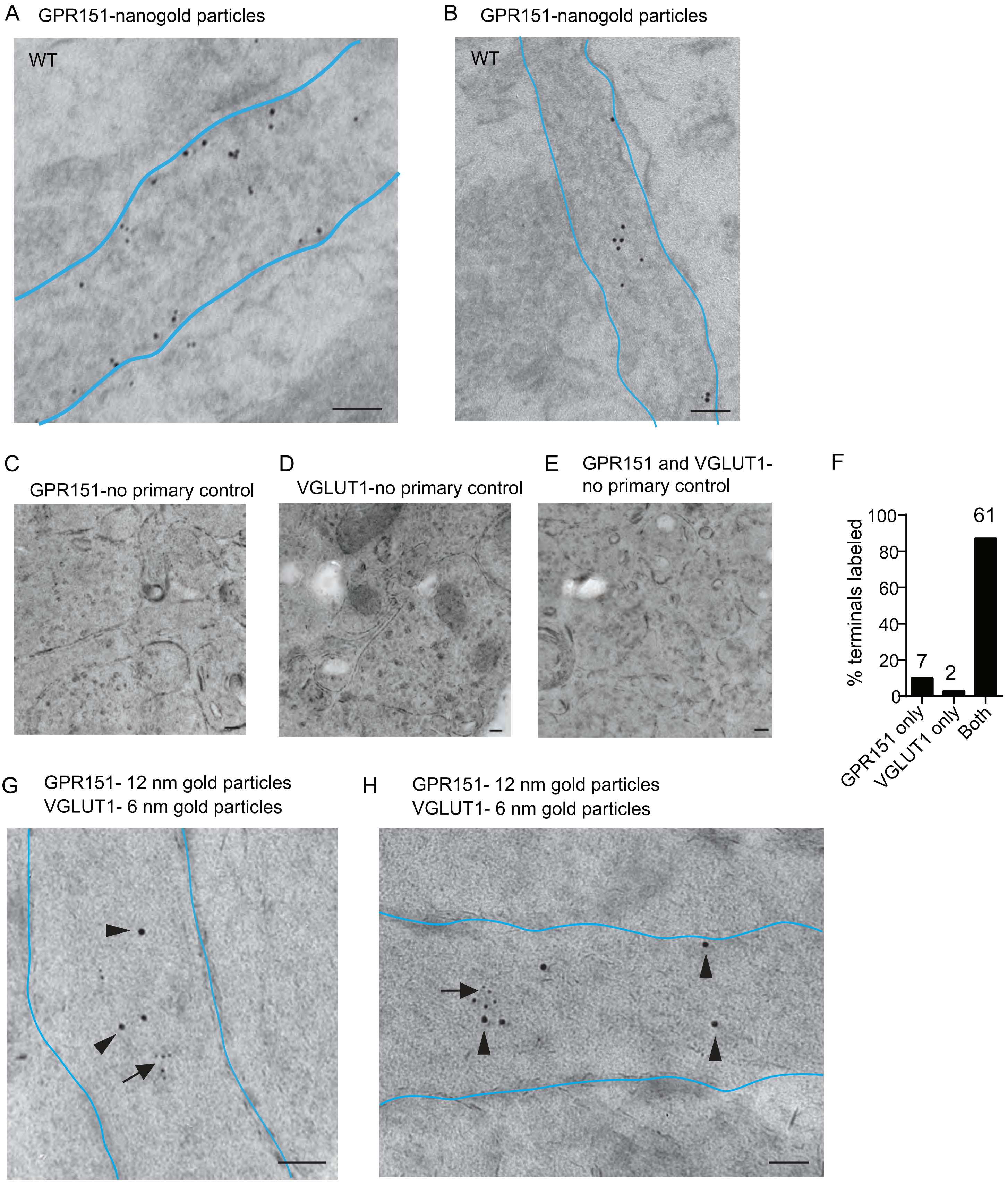

### supplemental Fig 10

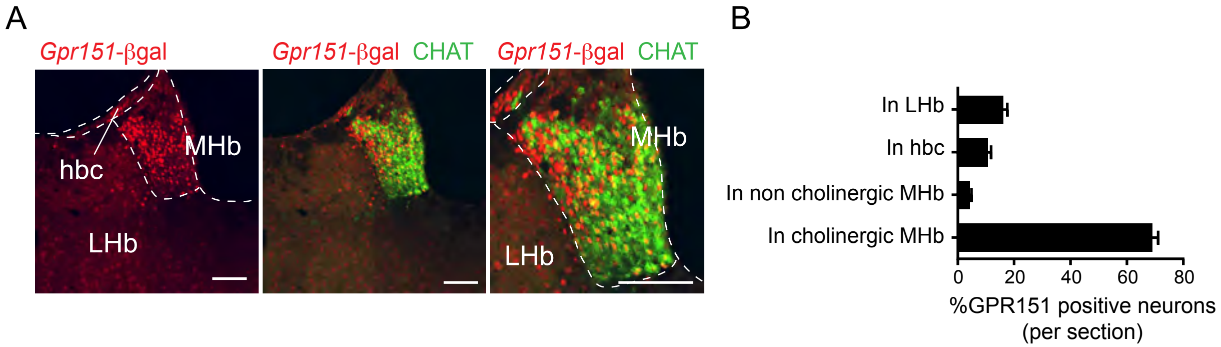

### supplemental Fig 11

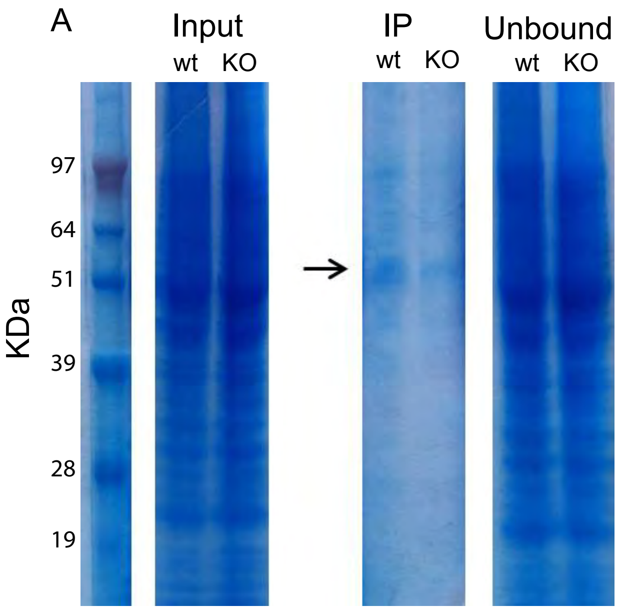

### supplemental Fig 12

A

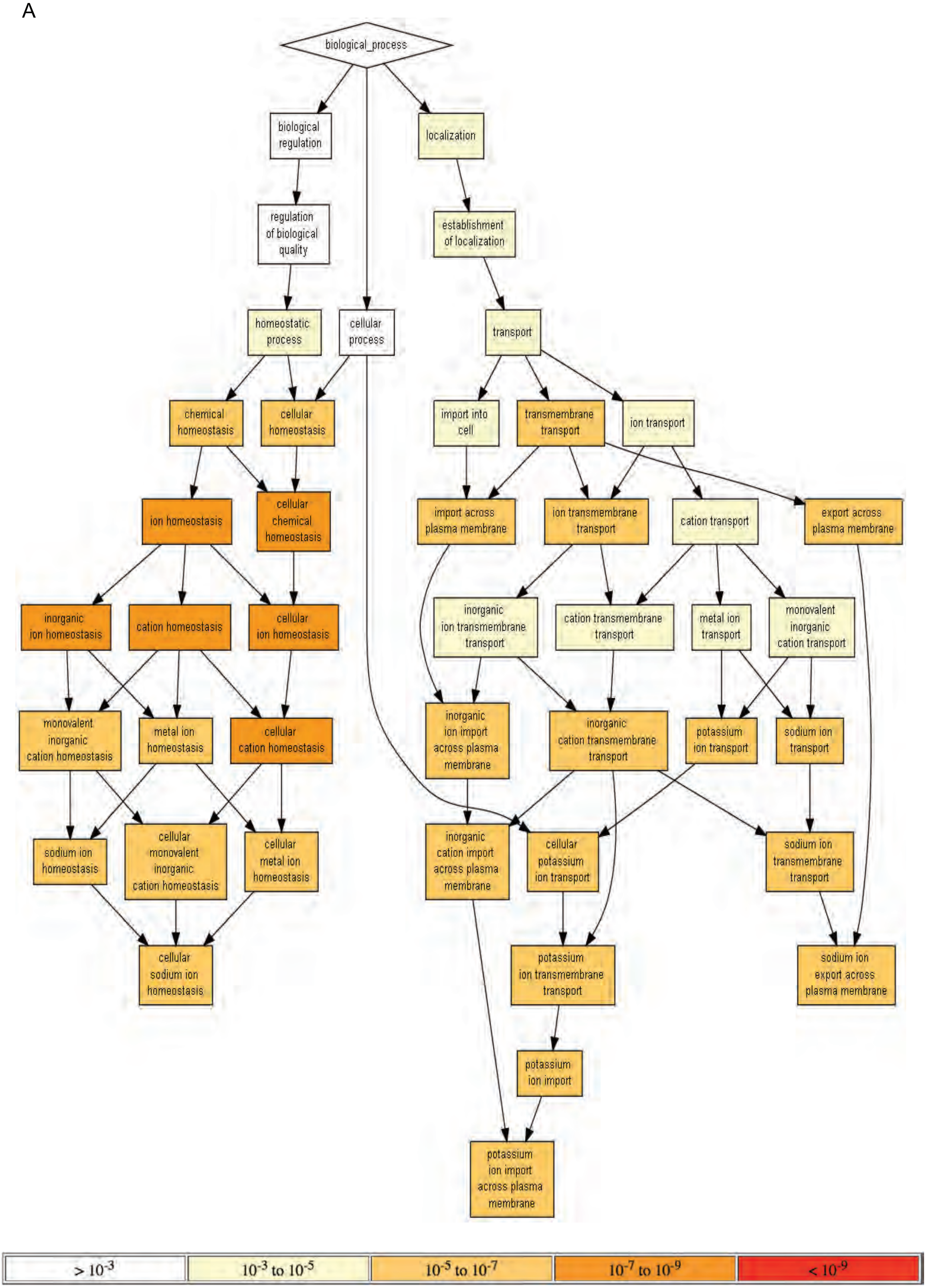


P-value color scale

### supplemental Fig 13

A

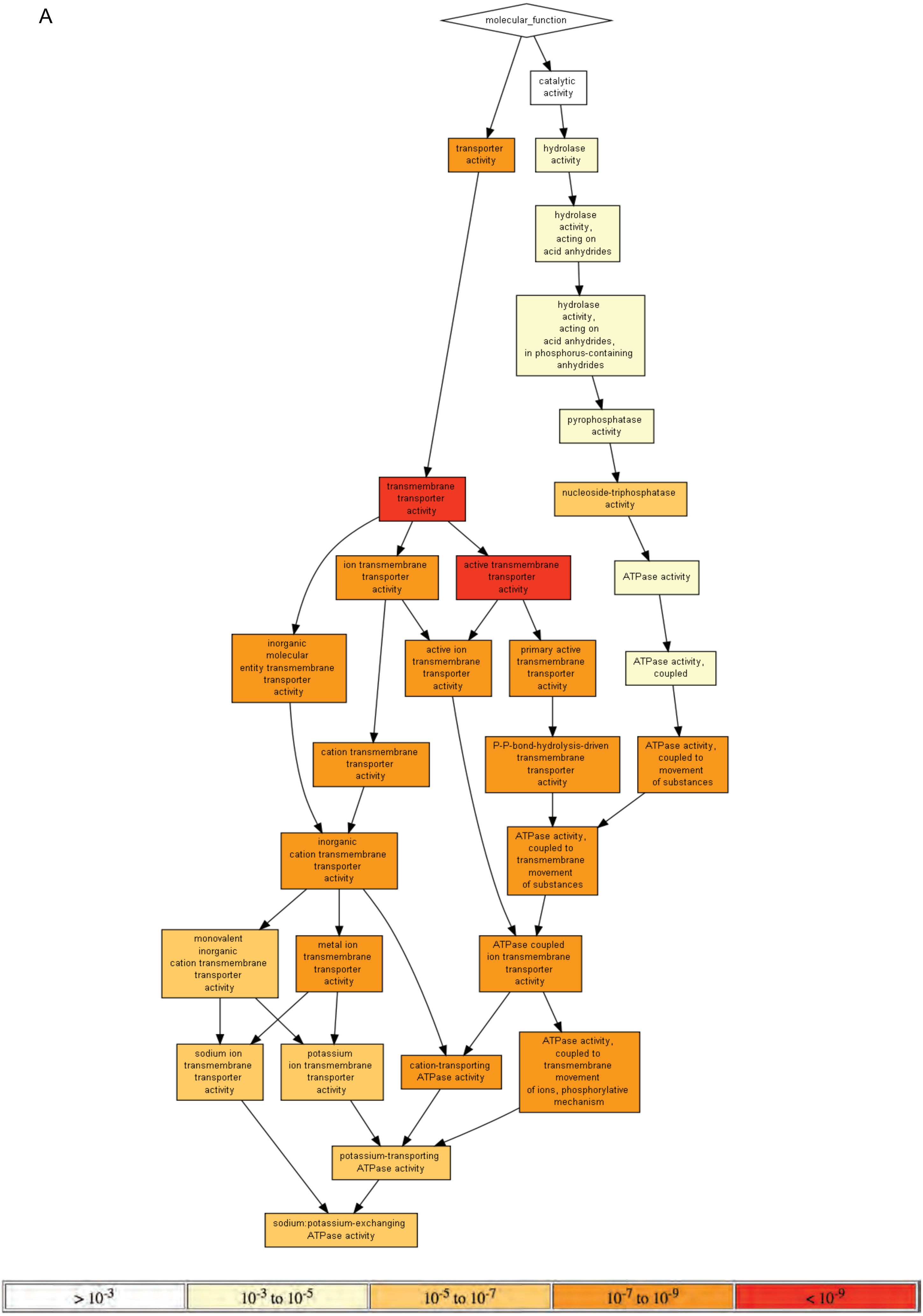


P-value color scale

### supplemental Fig 14

A

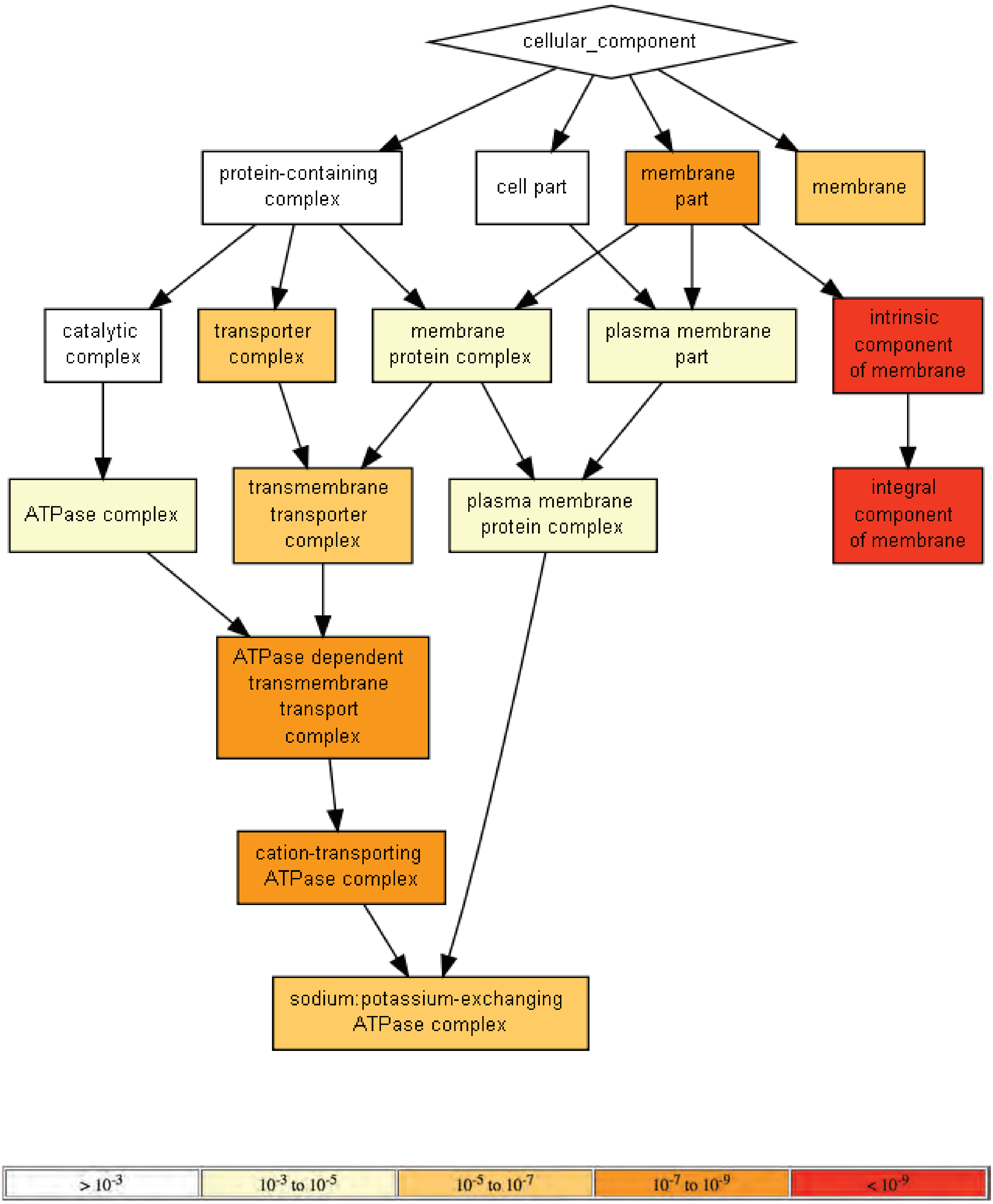


P-value color scale

### supplemental Fig 15

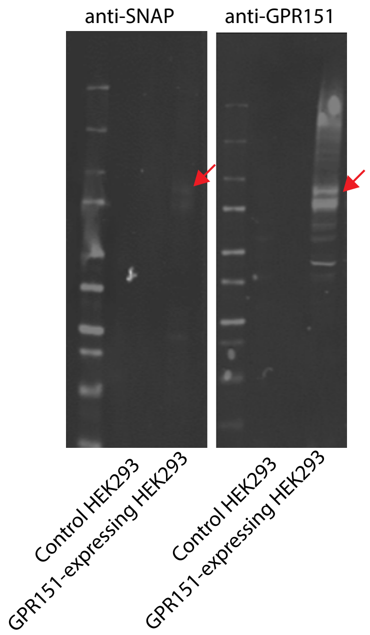
