## supplemental Figure legends and methods for "The Habenular Receptor GPR151 Regulates Addiction Vulnerability Across Drug Classes"

**Figure S1: Related to Figure 1**

(A) Integrative genome viewer (IGV) of TRAP data collected from the MHb of *Chat-EGFP-L10a* mice crossed to WT or *Gpr151-*KO showing that *Gpr151* is truncated at the beginning of the gene by the lacZ insertion to generate *Gpr151*-KO.

(B) Heat map of the top 50 genes in WT and KO MHb IP samples. *Gpr151* is indicated with an arrow.

**Figure S2: Related to Figure 1**

(A) Scatter plot comparing TRAP normalized read counts of MHb Immunoprecipitated (IP) samples from *Chat-EGFP-L10a* control mice (WT) to IP samples from *Chat-EGFP-L10a* x*Gpr151*-KO mice. Upregulated genes in *Gpr151*-KO are shown in pink, downregulated genes in blue (adjusted p value < 0.05, log2 fold change cutoff = 0.6 (absolute value)) (n=6 biological replicates, each replicate contains the habenulae of 5 *Chat-EGFP-L10a* transgenic mice).

(B) Volcano plot of TRAP data collected from the MHb of *Chat-EGFP-L10a* x WT vs *Chat-EGFP-L10a* x *Gpr151-*KO.

(C) M (log ratio) A (mean average) plot of TRAP data collected from the MHb of *Chat-EGFP-L10a* vs *Chat-EGFP-L10a* x *Gpr151-*KO.

**Figure S3. Related to Figure 1**

(A) Schematic diagram of the habenula-IPN pathway. MHb neurons project along the fasciculus retroflexus to the IPN, where they terminate in a zig-zag pattern.

(B-D) Coronal sections of the habenula showing double immunostaining of (B) GPR151 (red) and nAChRα3-GFP (green) in Tabac mice expressing the *Chrnβ4-α3-eGFP-α5* gene cluster; (C) GPR151 (red) and the µ-opiod receptor (MOR) (green) in WT mice; (D) GPR151 (red) and the cannabinoid receptor 1 (CB1) (green) in WT mice; The Manders’ colocalization index (M1) and the significance of correlation (P) were measured on the high magnification picture of the indicated square area. Scale bar: 100 µm for low magnification pictures, 20 µm for high magnification pictures. Colocalization of GPR151 and nAChRα3-GFP and GPR151 and MOR was observed in the lateral part of MHb. Very little overlap was observed between GPR151 and CB1R on the central part of the MHb.

(E-J) Coronal sections of the IPN showing double immunostaining of (E, F) GPR151 and nAChRα3-GFP in Tabac mice expressing the *Chrnβ4-α3-eGFP-α5* gene cluster, (G,H) GPR151 and MOR and (I,J) GPR151 and CB1R. The Manders’ colocalization index (M1) and the significance of correlation (P) were measured on the high magnification pictures. Scale bar: 100 µm for low magnification pictures, 20 µm for high magnification pictures of E,G,I, 10 µm for F,H,J.

**Figure S4. Related to Figure 1.**

(A) Coronal sections of the IPN showing that MOR immunostaining is similar in WT and *Gpr151*-KO mice. Scale bar: 100 µm.

(B) Quantification of MOR immunofluorescence signal shows no significant differences between WT and *Gpr151*-KO in both habenula and IPN (WT habenula: 100 ± 8.38, n=4; KO habenula: 109.2 ±4.04, n=5, unpaired t-test, p=0.32. WT IPN: 100 ± 2.76, n=5, KO IPN: 108.7 ± 6.50, n=5, unpaired t-test, p=0.25).

(C) Coronal sections of the IPN showing that CBR1 immunostaining is similar in WT and *Gpr151*-KO mice. Scale bar: 100 µm.

(D) Quantification of CBR1 immunofluorescence signal shows no significant differences between WT and *Gpr151*-KO in both habenula and IPN (WT habenula: 100 ± 26.77, n=4; KO habenula: 97.94 ± 22.95, n=4, unpaired t-test, p=0.95. WT IPN: 100 ± 17.24, n=6, KO IPN: 113.6 ± 17.37, n=4, unpaired t-test, p=0.61).

Data are represented as mean ± SEM.

**Figure S5. Related to Figure 2**

(A) Basal locomotor activity, measured as number of beam breaks, was not significantly different between WT and *Gpr151*-KO mice (n=8-12, repeated measures 2-way ANOVA).

(B) Anxiety-like behavior, measured in the elevated plus maze, was not significantly different in *Gpr151*-KO compared to WT mice. The time spent in the closed and open arms was similar between genotypes (Closed arms: WT=468.8 ± 17.10, n=8; KO=442.3 ± 10.91, n=12, unpaired t-test, p=0.18. Open arms: WT=44.49 ± 9.48, n=8, KO= 42.37 ± 6.94, n=12, unpaired t-test p=0.85).

(C) Anhedonia-like measured with the sucrose preference test was not different between WT and *Gpr151*-KO mice (WT=94.11 ± 0.20, n=10; KO=91.91 ± 1.78, n=10, unpaired t-test, p=0.23).

(D) *Gpr151*-KO mice exhibited normal percent prepulse inhibition of the acoustic startle response across prepulse (PP) intensities (3, 6, 12 dB above background) (n=8-12, 2-way ANOVA).

(E) Total locomotor activity during 30 min after injection of saline or of the CB1R agonist ACEA (5mg/kg, i.p). Injection of ACEA decreased locomotor activity in both WT and *Gpr151*-KO mice (n=9 per genotype; repeated measures 2-way ANOVA, Sidak’s multiple comparisons test ***p<0.001 saline vs ACEA in both WT and KO).

(F) Quantification of the locomotor activity for 30 min, measured as number of beam breaks, immediately after injection of ACEA (5 mg/kg, i.p.). No differences were observed between genotype (n=9 per genotype; repeated measures 2-way ANOVA).

(G-H) Food intake measurements of WT (G) and *Gpr151*-KO (H) mice during 1 hr after an injection of saline on the first day and an injection of the CB1R agonist, ACEA (5mg/kg, i.p.) on the second day. Each dot represents one mouse (n=8-7 per genotype, paired t-test, WT p=0.0025, *Gpr151*-KO p=0.95).

(I) Body weight of WT and *Gpr151*-KO mice over age on regular water and food. No differences were observed in the non-linear regression curve between WT and *Gpr151*-KO (WT n= 61 different mice weighted at 15 different ages, *Gpr151*-KO n= 67 different mice weighted at 13 different ages).

Data are represented as mean ± SEM. See Table S3 for details of statistical analysis.

**Figure S6. Related to Figure 2**

**(**A) Schematic representation of the lentiviral contruct encoding *Gpr151*-HA-tagged and the lentiviral bipromoter construct expressing EGFP and shRNA against *Gpr151*.

(B) Western blot analysis of non-transduced HEK293Tcells (line1), transduced with HA-*Gpr151* alone (line 2) or together with 5 different shRNAs against *Gpr151* (lines 3-7). GPR151 was detected in line 2 (47 kDa: GPR151) and was not observed in line 5, indicating that shRNA3 was efficient at knocking-down *Gpr151* in transduced cells.

(C) GPR151 immunostaining of habenula and IPN coronal sections of *Gpr151*-KO mice injected in the MHb with AAV2/1-*Gpr151* (to rescue expression) or AAV2/1-control virus. Scale bar: 100um.

**Figure S7. Related to Figure 3**

(A) Active (solid line) and inactive (dotted line) lever presses during food-training (n=12 WT, 14 KO).

(B) Active (solid line) and inactive (dotted line) lever presses during nicotine self-administration. n= 6 WT, 6 KO (Mixed-effects model, Tukey’s multiple comparison test).

Data are represented as mean ± SEM. See Table S3 for details of statistical analysis.

**Figure S8. Related to Figure 4**

(A-B) Representative micrographs of pre-embedding immunogold EM analyses showing GPR151 immunogold particles at habenular terminals of WT mice indicating synaptic vesicles (SV) close to the active zone (A) or without the active zone on the plane of the picture (B). The area of the presynaptic terminal is delineated in blue. GPR151 nanogold particles were mainly located at the plasma membrane. Scale bar: 500 nm. (DCV, dense core vesicle, mit, mitochondria).

(C-D) Representative micrographs of pre-embedding immunogold EM analyses showing the absence of GPR151 nanogold particles in presynaptic terminals of *Gpr151*-KO mice. Scale bar: 500 nm.

(E) Percentage of synaptic contacts labeled with GPR151 that are axo-dendritic (50/53), axo-axonic (2/53) and axo-somatic (1/53).

(F-G) Frequency histograms showing the percentage of GPR151 nanogold particles labeling either a SV (F) or a DCV (G) located at the indicated distances (100 nm bins) from the central point of the AZ (n=112 GPR151 particles labeling SV and n=11 GPR151 particles labeling DCV measured in n=50 terminals).

(H) Quantitative analysis of GPR151 pre-embedding immunogold electron micrographs of synaptic terminals with SV, without the active zone on that plane. The percentage of GPR151 nanogold particles per terminal located on the membrane is 85.87% ± 2.14; in association with SV is 5.7% ± 1.5; in association with DCV is 0.6% ± 0.34; and in other unidentified structures is 7.81 ± 1.31 (n=46 terminals, total of 864 particles).

Data are represented as mean ± SEM.

**Figure S9. Related to Figure 4.**

(A-B) Transversal sections of habenular axons showing GPR151 immunogold particles at the membrane (A) and inside the axon along the microtubules being transported towards the terminal (B). Scale bar: 100 nm.

(C-E) Control experiments of single and double post-embedding labeling of GPR151 and VGLUT1 in the IPR of WT mice. Sections were treated following the same procedure while omitting the primary antibody. Scale bar: 100 nm.

(F) Quantification of the double-postembedding labeled terminals showing the number of terminals labeled only with GPR151, VGLUT1 or both and corresponding percentages.

(G-H) Transversal sections of habenular axons showing GPR151 (12 nm) and VGLUT1 (6 nm) particles at higher magnification. Scale bar: 100 nm.

Data are represented as mean ± SEM.

**Figure S10. Related to Figure 5**

(A) *Gpr151*-KO mice were generated by insertion of the the β-galactosidase (β-Gal) reporter under the *Gpr151* promoter (see Figure S1A).. Coronal section of a *Gpr151*-KO mouse, showing β-Gal and CHAT immunostaining in the habenula. The right panel is a high magnification of the medial habenula (MHb). Scale bar: 100 µm.

(B) Quantification of β-Gal positive neurons in the cholinergic part of MHb (69.06 % ± 2.02); in the non-cholinergic part of MHb (4.30 % ± 0.56), in the habenular commissure (hbc) (10.51 % ± 1.31) and in the lateral habenula (LHb) (16.14 % ± 1.44) (n=39 habenulae from 3 different mice).

Data are represented as mean ± SEM.

**Figure S11. Related to Figure 6**

(A) Commassie-stained gels of brain samples after GPR151 immunoprecipitation showing input samples, IP samples and flow-through samples of WT and *Gpr151*-KO IPN samples. A double band corresponding to GPR151 (around 53KDa, arrow) was observed in the IP sample of WT but not in *Gpr151*-KO.mice.

**Figure S12. Related to Figure 6**

(A) Gene ontology analysis showing the biological processes of the proteins co-immunoprecipitated with GPR151 in WT mice identified by mass spectrometry (See also Table S4).

**Figure S13. Related to Figure 6**

(A) Gene ontology analysis showing the molecular functions of the proteins co-immunoprecipitated with GPR151 in WT mice identified by mass spectrometry (See also Table S4).

**Figure S14. Related to Figure 6**

(A) Gene ontology analysis showing the cellular component of the proteins co-immunoprecipitated with GPR151 in WT mice identified by mass spectrometry (See also Table S4).

**Figure S15. Related to Figure 7**

(A) Western blots of HEK293 cells untransfected (parental cell line) and stably transfected with 5HT3SigP-SNAP-GPR151 (GPR151-HEK293 stable clonal cell line). Left panel shows anti-SNAP antibody staining (arrow). Right panel shows anti-GPR151 antibody signal in the stable clonal cell line. The theoretical molecular weight for GPR151 is 46 and 53 kDa, for 5HT3SigP-SNAP-GPR151 is around 66kD, while the western blot results show a molecular weight between 70-100kD likely due to glycosylation.

**SUPPLEMENTAL EXPERIMENTAL PROCEDURES**

**Animals**

Transgenic *Chat-EGFP-L10a* (*Chat-DW167*) have been previously described (Doyle et al., 2008; Frahm et al., 2011; Gorlich et al., 2013; Heiman et al., 2008). Tabac mice expressing the *Chrnβ4-α3-eGFP-α5* gene cluster have been characterized in (Frahm et al., 2011). *Gpr151* knockout mice ([Gpr151tm1Dgen](http://www.informatics.jax.org/accession/MGI:3604578" \t "_top)) were obtained from Deltagen (RRID:MGI_3606630). They were backcrossed to C57BL/6 for eight generations. *Chat-*ChR2-YFP BAC transgenic mice were obtained from The Jackson Laboratory (Strain name: B6.Cg-Tg(Chat-COP4*H134R/EYFP)6Gfng/J, RRID:MGI_5491602). All lines were maintained on a heterozygous background.

**Drugs**

(-) Nicotine hydrogen tartrate salt was purchased from Sigma-Aldrich (St. Louis, MO, USA). Nicotine concentrations refer to the free base. For nicotine self-administration experiments, (-) nicotine hydrogen tartrate salt was dissolved in 0.9% sterile saline for 0.3 mg/ml stock solution. For systemic injections, nicotine was dissolved in sterile water and injected at 10 ml/kg volume. The pH of all solutions was adjusted to approximately 7.4. Morphine sulfate salt pentahydrate was obtained from Sigma-Aldrich (St. Louis, MO, USA) and dissolved in sterile water. Arachidonyl-2'-chloroethylamide (ACEA) was obtained from Tocris Bioscience (UK) pre-dissolved in anhydrous ethanol (5 mg/ml). Ethanol was evaporated and ACEA was re-dissolved in dimethyl sulfoxide (DMSO) the day of the experiment. The same amount of DMSO was used for the control group. The final amount of DMSO in saline was 0.04%. Drugs were dissolved in 0.9 % saline and administered in a volume of 100 µl per 10 g body weight. Isobutylmethylxanthine (IMBX) and forskolin were obtained from Tocris Bioscience and Sigma-Aldrich.

**Immunohistochemistry of mouse brain samples and quantification analysis**

The primary antibodies used were: rabbit polyclonal anti-GPR151 (1:1000, Sigma, RRID:AB_10743863), goat polyclonal anti-GPR151 (1:500, Santa Cruz, RRID:AB_2113821), goat polyclonal anti-CHAT (1:1000, Millipore, RRID:AB_2079751), chicken polyclonal anti-ß-galactosidase (β-Gal) (1:200, Abcam, RRID:AB_307210), chicken polyclonal anti-GFP (1:1000, Aves, RRID:AB_10000240), rabbit polyclonal anti-mu opioid receptor (1:1000, Immunostar, RRID:AB_10730727), rabbit polyclonal anti-cannabinoid receptor 1 (1:1000, Abcam, RRID:AB_447623). The sections were incubated with primary antibodies overnight at 4°C or room temperature. After incubation with secondary antibodies (Jackson ImmunoResearch, PA, USA), sections were washed, mounted on slides and coverslipped in immu-mount (Thermo Scientific, MA, USA). Tyramide signal amplification (PerkinElmer, MA, USA) was performed for the cannabinoid receptor 1 staining to enhance the signal. Heat-mediated antigen retrieval, 15 minutes at 95°C in citric acid (pH 6.0), was performed prior to incubation with the choline acetyltransferase (CHAT) antibody. Fluorescent signals were detected using a confocal laser scanning microscope (Zeiss LSM700, Germany). Image J was used for quantification analysis. The number GPR151 (β-Gal positive) and CHAT cells per section was quantified in 39 habenula of 3 different GPR151-KO mice.

Colocalization analysis was done as described in (Frahm et al., 2015) with the Coloc 2 plugin in the Fiji image processing package. Background was eliminated by median subtraction (Dunn et al., 2011). Manders' colocalization coefficients (M1 and M2), which are proportional to the number of colocalizing pixels in each channel relative to the total number of pixels, were calculated (Manders et al., 1993). M1 or M2 > 0.55 indicates colocalization (Zinchuk and Grossenbacher-Zinchuk, 2014). In this study, only M1 is presented and refers to the proportion of pixels with GPR151 immunoreactivity that colocalize with the second marker. Costes' test for statistical significance was used to determine that the colocalization coefficients obtained were not due to random effects (Costes et al., 2004).  This test creates random images by shuffling blocks of pixels of one channel, measuring the correlation of this channel with the other (unscrambled) channel of the same image. The test was performed 100 times per image and the resulted P value indicates the proportion of random images that have better correlation than the real image. A P-value of 1.00 means that none of the randomized images had better correlation.

**Co-immunoprecipitation and western blot**

The IPN was dissected from 15-20 WT and *Gpr151*-KO mice and homogenized in 500 ul lysis buffer (50 mM Tris-HCl pH 8.0, 100 mM NaCl, 2mM EDTA, 1% Triton-X100, 5% Glycerol and protease and phosphatase inhibitors tablets (Roche) using a tissue grinder. The homogenates were placed on a rotator for 30 min at 4 °C centrifuged for 15 min at 14000 rpm. The supernatant, which contains the cytosolic proteins, was collected and the pellet, which contains the membrane protein, was resuspended in 50-100 ul lysis buffer containing 2% Triton X-100 for 2 hr at 4°C. Samples were centrifuged for 15 min at 14000 rpm and the supernatant was combined with the cytosolic sample.

For co-immunoprecipitation for mass spectrometry analysis, M-270 Epoxy Dynabeads (Invitrogen, MA, USA) were used. Coupling of the antibody to the beads was done as suggested by the manufacturer. A combination of three different GPR151 antibodies (30 µg of rabbit polyclonal anti-GPR151 (Sigma, RRID:AB_10743863), 10 µg of goat polyclonal anti-GPR151 (Santa Cruz, RRID:AB_2113821) and 20 µl of mouse polyclonal anti-GPR151 (Sigma, RRID:AB_10608138) were coupled to 15 mg of beads per sample during 24 hr at 37°C in 1.5 ml volume. After conjugation, beads were washed, added to the sample and incubated for 16 hr on a rotator at 4°C. The beads were subsequently washed six times with lysis buffer (twice with buffer containing Triton-X100 and three additional times with lysis buffer without Triton-X100). The suspension was then transferred to a clean tube and two serial elutions of the protein complex were performed with 40 ul 8M Urea (freshly prepared) for 30 and 10 min at room temperature on a rotator. The supernatant (80 µl total) was transferred to a new tube and 500 ul of cold acetone was added to precipitate proteins and remove detergent. The sample was kept overnight at -20 °C and centrifuged the following morning at 14000 rpm for 5 min. The acetone was discarded and the pellet was dissolved in 8M urea for mass spectrometry analysis.

For co-immunoprecipitation for western blot analysis, 50 µl of DynaBeads Protein G (Invitrogen) were used per sample according to the manufacturer’s recommendations. 5 µg of rabbit polyclonal anti-GPR151 (Sigma, RRID:AB_10743863) and 1 µg of goat polyclonal anti-GPR151 (Santa Cruz, RRID:AB_2113821) were incubated with the beads for 1 hr at room temperature. After washing the beads and antibody complex, the sample was added to the tube and incubated with the beads for 16 hr on a rotator at 4°C. The complex was eluted with 20 µl elution buffer plus 10 µl of NuPAGE LDS sample buffer/reducing agent mix and incubated 10 min at 70 °C. The supernatant was removed and directly loaded onto a NuPAGE Bis–Tris 4–12 % pre-cast gel (100 V, 2 hr). Proteins were transferred onto a PVDF membrane for 90 min at 100 V in the NuPAGE transfer buffer supplemented with 20 % methanol. The membranes were then blocked for 2 h in 5 % BSA in TBS-T and incubated with mouse polyclonal anti-GPR151 (1:1000, Sigma, RRID:AB_10608138), mouse monoclonal anti-Gαo 1/2 (1:1000, Synaptic Systems, #271 111) and rabbit polyclonal anti-Gαs (1:1000, Abcam, RRID:AB_1860300) overnight at 4°C. LI-COR secondary antibodies were used (1:15000) for 1 hr and the membranes were scanned and quantified using the LI-COR Odyssey imaging system (LI-COR Biosciences, NE, USA). Comassie stainings were performed using Coomassie® R-250 (ThermoFisher Scientific) following the manufacturer’s instructions.

**Proteomics Methods and data Analysis**

Ten samples (five biological replicates of each WT and *Gpr*151-KO immunoprecipitation experiments) were denatured in 8M urea, reduced with 10 mM DTT, and alkylated with 50 mM iodoacetamide. This was followed by proteolytic digestion with endoproteinase LysC (Wako Chemicals, VA, USA) overnight, and with trypsin (Promega, WI, USA) for 6h at room temperature. The digestion was quenched with 5% formic acid (final concentration) and resulting peptide mixtures were desalted using in-house made C18 Empore (3M) StAGE tips. Samples were dried and resolubilized in 2% acetonitrile and 2% formic acid. One fifth of each desalted (Rappsilber et al., 2007) sample was injected for analysis by reversed phase nano-LC-MS/MS (Ultimate 3000 coupled to a QExactive Plus, Thermo Scientific, MA, USA). After loading on a C18 PepMap trap column (5 µm particles, 100µm x 2 cm, Thermo Scientific, MA, USA) at a flow rate of 3 μl/min, peptides were separated using a 12 cm x 75µm C18 column (3 µm particles, Nikkyo Technos Co., Ltd. Japan) at a flow rate of 200 nL/min, with a gradient increasing from 5% Buffer B (0.1% formic acid in acetonitrile) / 95% Buffer A (0.1% formic acid) to 40% Buffer B / 60% Buffer A, over 140 minutes. All LC-MS/MS experiments were performed in data dependent mode with lock mass of m/z 445.12003. Precursor mass spectra were recorded in a 300-1400 m/z range at 70,000 resolution, and fragment ions at 17,500 resolution (lowest mass: m/z 100) in profile mode. Up to twenty precursors per cycle were selected for fragmentation and dynamic exclusion was set to 60 seconds. Normalized collision energy was set to 27.

Data were quantified and searched against Uniprot mouse database (February 2019) using MaxQuant (version 1.6.0.13) (Cox et al., 2011). Oxidation of methionine and protein N-terminal acetylation were allowed as variable modifications, cysteine carbamidomethyl was set as a fixed modification and two missed cleavages were allowed. The “match between runs” option was enabled, and false discovery rates for proteins and peptide spectrum matches were set to 1% and 2%, respectively. Protein abundances were expressed as intensity Based Absolute Quantitation (iBAQ) (Schwanhausser et al., 2011). In short: LOG2 transformed iBAQ signals were normalized by subtracting median. Identified proteins were further filtered by requiring that at least 3 signals were present for each protein in at least one condition. Missing signals were imputed and the two conditions were compared using a P-value only t-test. Threshold of P>0.05 and a difference of 2 linear folds were set as filters to identify proteins of interest.

**Immunohistochemistry and protein extraction of human brain samples**

Fixed samples of about 2 cm wide x 5 cm long x 1 cm deep containing the habenula, the fasciculus retroflexus and the interpeduncular nucleus, were cryoprotected for 2 days in sucrose in 15% sucrose and for 2 following days in 30% sucrose. 50 µM sections were sectioned on a SM2000R sliding microtome (Leica, IL, USA). Sections were washed for one hour in 1xPBS containing 0.2%TX100 and then blocked for 2 hours in 1xPBS containing 0.5%TX100 and 4%NDS. Subsequently, sections were incubated overnight at 4 °C with the rabbit polyclonal anti-GPR151 (Sigma, 1:1000, RRID:AB_10743863) diluted in blocking buffer. The following morning, sections were washed with 1xPBS containing 0.2%TX100 and incubated for 2 h at room temperature with secondary antibodies (1:500; Jackson ImmunoResearch, PA, USA). Sections were mounted on slides and examined and photographed with a LSM700 (Zeiss, Germany) confocal microscope.

For human western blot analysis, the IPN (60-450 mg) was dissected from frozen sections of the midbrain and homogenized using a motor-driven Teflon-glass homogenizer in 500 µl - 1 ml lysis buffer. Cytosolic and membrane protein fractions were used for immunoprecipitation and western blot analysis as described with mouse tissue.

**Stereotaxic viral injections**

Mice, 8-9 weeks old, were anesthesized with ketamine/xylazine (130 mg/kg and 10 mg/kg, respectively), and placed in a Benchmark stereotaxic frame with a Cunningham mouse adaptor (Leica). The eyes were covered with Regephitel® ointment to prevent desiccation. Four holes were drilled on the skull with a micromotor drill with a 0.35 mm diameter bit. The coordinates used to target the medial habenula were the following: anterior-posterior (from Bregma): -1.4 and -1.75; lateral +/-0.33; dorso-ventral (from the skull) -2.72 and -2.70. Glass PCR micropipets (Drummond) that had been pulled with a microelectrode puller (Narashige) to create a narrow, fine tip of ~10-12 mm length were used for the injections. The cannula was filled from the backside with virus, followed by mineral oil. A metal plunger was inserted into the upper part of the cannula and placed in the stereotaxic apparatus. 0.5-1 µl of virus was injected into the mouse medial habenula using a MO-10 oil hydraulic micromanipulator (Narashige) with a flow rate of ~0.1 µl/min. The cannula was left in place for 5 min, allowing the virus solution to disperse in the tissue, before it was slowly retracted. The incision was closed with the tissue adhesive Vetbond (3M). After surgery, mice were warmed up by the infrared lamp and left for recovery for 2h. Mice were kept under S2 conditions for a week. The following virus were used: AAV2/1-U6-Gpr151-shRNA and AAV2/1-GFP-U6-scramble-shRNA from Vector Biolabs, AAV2/1-CAG-M-Gpr151-WPRE from Vector Biolabs for rescue experiments, , AAV2/1-L10a-EGFP control virus from Janelia farms. Viral titers were 10^13^ inclusion forming units (IFU)/mL.

**Electron microscopy**

Fixation, high pressure freezing and freeze substitution was performed as in (Frahm et al., 2015). For pre-embedding nanogold immunolabeling, IPN coronal sections from WT and *Gpr151*-KO mice were incubated in blocking solution (3% BSA, 0.1% saponin) for 2 hours at room temperature. The sections were then single labeled at 4°C for 36 hr with rabbit polyclonal anti-GPR151; 1/3000 (Sigma, RRID:AB_10743863) diluted in blocking solution and incubated in secondary antibody diluted 1:100 for 1hr at room temperature (Nanoprobes- Nanogold, anti-Rabbit: 2003). The sections were fixed in 2.5% glutaraldehyde overnight at 4°C and subsequently underwent silver enhancement (HQ Silver Enhancement 2012, Nanoprobes, NY, USA), and Gold Toning using a 0.1% solution of gold chloride (HT1004, Sigma-Aldrich, MO, USA). The rostral IPN was excised and postfixed with 1% osmium tetroxide in the buffer for 1 hour on ice. Sections underwent *en bloc* staining with 1% uranyl acetate for 30 min, dehydration in a graded series of ethanol, 10 minutes in acetone, infiltration with Eponate 12™ Embedding Kit (Ted Pella, CA, USA), embedding  with the resin and polymerization for 48h at 60°C.  70nm ultrathin sections were analyzed on a JEOL JEM-100CX (JEOL, Tokyo, Japan) at 80kV using with the digital imaging system (XR41-C, Advanced Microscopy Technology Corp, Woburn, MA, USA).

Post-embedding immunolabeling was carried out on 90 nm ultrathin sections mounted on carbon film 200 mesh nickel grids (Electron Microscopy Sciences, PA, USA). Sections were incubated in blocking solution containing 3% BSA, 0.1% in 1xPBS for 1h, incubated with single or double primary antibodies: rabbit polyclonal anti-GPR151; 1/800 (Sigma, RRID:AB_10743863) and/or guinea pig polyclonal anti-VGLUT1; 1/400 (Synaptic Systems, RRID:AB_887878) diluted in blocking solution overnight at 4°C, rinsed in blocking solution for 30 min, incubated with appropriate secondary antibodies (e.g., anti-rabbit IgG conjugated to 12 nm gold particles and anti-guinea pig IgG conjugated to 6 nm gold particles (Jackson ImmunoResearch, PA, USA) diluted 1/20 in PBS containing 0.5% BSA and 0.1% saponin for 1h at room temperature. Finally sections were washed twice in PBS and twice in ultrapure water, stained with1% osmium tetroxide and lead citrate and examined in a JEOL JEM-100CX microscope. The control experiment was done by following the same procedure except for omitting the primary antibody and incubating with blocking solution instead.

Quantitative analyses were performed on preembedding nanogold labeled synaptic terminals (at 10000x magnification) from three WT mice and *Gpr151*-KO mice. 45 terminals with distinguishable active zone and synaptic vesicles of WT and *Gpr151*-KO mice were analyzed to quantify the area of the terminal, length of the active zone, and size, density and distribution of synaptic vesicles. On a separate analysis, 50 terminals of WT mice with a distinguishable active zone and 46 terminals filled with vesicles but without the active zone in the picture were quantified (a total of 999 GPR151 nanogold particles in terminals with AZ and 864 particles in terminals without AZ). All measurements were done with ImageJ.

**TRAP and RNA-SEQ**

RNA-seq reads were aligned to the UCSC mm10 reference genome using STAR (Dobin et al., 2013), version 2.3.0e_r291, with default settings. Quantification of aligned reads was done using htseq-count module, part of the ‘HTSeq’ framework (Anders et al., 2015), version 0.6.0, using default settings, with ‘union’ mode to handle reads overlapping more than one feature. Differentially expressed genes were identified by performing a negative binomial test using DESeq2 (Love et al., 2014) (R-package version 1.4.5) with default settings. Significant P-values were corrected to control the false discovery rate of multiple testing according to the Benjamini-Hochberg (Benjamini and Hochberg, 1995) procedure at 0.05 threshold. Gene lists were then further filtered for basemean > 100 counts to exclude genes with low expression.

**Behavioral analysis**

Open field. Locomotor activity was measured in eight identical open field boxes (50x50x22.5 cm) equipped with two rows of infrared photocells placed 20 and 50 mm above the floor (Accuscan & Omnitech Electronics, OH, USA). Mice were placed in the center of the field and activity was recorded by the Fusion Software. To habituate the animals to the test environment and to obtain a stable baseline, locomotor activity was measured immediately after a saline injection for 60 min 2 days before the testing day.

To evaluate the hypolocomotor effects of nicotine (0.65 mg/kg, i.p), mice were placed on open field boxes for 20 min immediately after injection. Tolerance to the locomotor effect of nicotine was measured every other day for 20 min immediately after nicotine injection for 11 days. The acute effects of morphine (10 mg/kg, s.c.) on locomotion were measured for 60 min immediately after injection. To measure sensitization to the locomotor effect of morphine, mice were given daily injections of morphine (10 mg/kg, s.c.) for 6 days and locomotor activity was monitored every other day for 60 min immediately after injection. The effect of ACEA (5mg/kg, i.p.) on locomotion was measured on open field boxes for 30 min immediately after injection.

Elevated Plus maze. Elevated Plus Maze was used to assess anxiety-like behavior. The apparatus consisted of a central platform (5 × 5 cm), two opposed open arms (25 × 5 cm) and two opposed closed arms with 15-cm-high black walls. The edges of the open arms were raised 0.25 cm to decrease the chance of a mouse falling. Mice were placed on the central platform and the time and activity were recorded for 10 min via a camera positioned above the apparatus. Time spent in each arm and the distance traveled was recorded automatically using EthoVision XT Tracking Software (Noldus, VA, USA).

Prepulse Inhibibition. Startle reflexes were measured in four identical startle response SR-LAB apparatus (San Diego Instruments, CA, USA). Each system contained a Plexiglas cylinder, 5.5 cm in diameter and 13 cm long, mounted on a platform located in a ventilated, sound-attenuated chamber. Startle responses were transduced by piezoelectric accelerometers mounted under each platform. Output signals were digitized and the average startle response and maximum amplitude of the startle response were recorded as startle units. The startle session began with a 5 min acclimation period in the presence of a 65-dB background white noise, followed by presentation of five 120 dB pulse trials to ensure a stable baseline and reduce variability. Afterwards mice were presented with four startle stimulus of different intensities (80, 90, 100, 110, 120 dB). A total of 20 trials in a pseudo-randomized manner with an average inter-trial-interval of 15 sec (range: 6-21 sec) were presented. Amplitude of startle was measured within a 100 msec window following the stimuli. Prepulse Inhibition was examined using prepulses of 3, 6 or 12 dB (20ms) above background. Testing consisted of twelve 120 dB pulses (40 ms duration) alone and ten pulses preceded (100 ms) by each prepulse (20 ms duration). One last block of five 120 db pulses was presented to ensure again a stable baseline. Percent PPI was calculated using the following formula: 100x(startle alone – startle with prepulse/startle alone). PPI was reported as percent inhibition of the average startle response.

Food intake, oral consumption and weight measurements. Animals were single housed one week before the experiment. They received sham injections of saline to get habituated to the procedure for two days before the testing day. On the testing day, mice were fasted for 2 hours before injection of saline or ACEA (5mg/kg, i.p.). The bedding of the cage was removed during experiment and pre-weighed food pellets (20 mg Dustless Precision Pellets, Bio-Serv, NJ, USA) were placed into the cage on a 6mm petri dish. Food intake was measured 1hr after injection.

To study the effects of ACEA on weight, male mice were injected with 5mg/kg ACEA at the same time every day for 7 days. Mice were weighted before and after the treatment.

To measure the effects of nicotine on weight, male mice were treated with nicotine (162,5 µg/ml) via the drinking water for 28 days. To minimize taste aversion, 2% saccharin was added to both, treatment and control groups.

Sucrose preference was measured for 24hr. Mice were single-housed and given a free choice between two bottles, one with 2% sucrose and another one with tap water. The solutions were presented in 50 ml falcon tubes containing stoppers fitted with ball–point sipper tubes to prevent leakage. To prevent possible effects of side preference, the position of the bottles was switched after 24 hr and the test was repeated. The preference for sucrose was calculated as a percentage of consumed sucrose solution of the total amount of liquid drunk.

**Operant conditioning**

Mice were mildly food restricted to 85-90% of free-feeding body weight and trained to respond on one of two levers (active and inactive) in an operant conditioning chamber (10 x 10 x 10 cm chamber in sound attenuating box, Med-Associates Inc., St. Albans, VT, USA) for food pellets (20 mg pellets; TestDiet, Richmond, IN). Mice responded during 1 h daily training sessions (7 sessions per week) until they achieved stable responding under a fixed-ratio 5, time out 20 sec (FR5TO20) schedule of reinforcement. Stable responding was defined as 3 consecutive sessions when mice achieved >25 food pellets/session. To test the effects of nicotine on food responding, mice were injected with saline or nicotine (1mg/kg, s.c.) 10 min prior to their regular daily session and effects on food responding were measured for 5 min. Nicotine and saline injections were delivered according to an unbiased counterbalanced design.

**Jugular catheter surgery**

Mice were anesthetized using isoflurane (1-3%)/oxygen vapor mixture and implanted with intravenous catheters. Catheters consisted of 6 cm (mice) length of silastic tubing fitted to a guide cannula (Plastics One, Wallingford, CT), curved to a right angle. Catheter/cannula assemble was encased in dental acrylic with 5 cm of catheter tubing extended from the cannula base. The catheter tubing was surgically impacted via an incision on the animal’s back, with the catheter passed subcutaneous to the neck. The catheter was then implanted approximately 1 cm into right jugular vein and secured in place using surgical silk suture. Catheters were flushed daily with physiological sterile saline solution (0.9% w/v) containing heparin (60 USP units/ml). Catheter integrity was tested with the acute reacting barbiturate anesthetic, Brevital (methohexital sodium, Eli Lilly, Indianapolis, IN).

**Intravenous nicotine self-administration procedure**

When mice demonstrated stable responding in the operant chamber (>25 food pellets per session), they underwent jugular catheter implantation. The animals were permitted at least 48 h to recover from surgery and then permitted to respond for food rewards again under the same FR5TO20 sec schedule. Once stable responding was re-established subjects were permitted to respond for intravenous nicotine infusions during 1 h daily sessions, 7 days per week (0.03 mg/kg/infusion). Nicotine was delivered via the implanted intravenous catheter by a Razel syringe pump (MedAssociates Inc., St Albans VT USA). Completion of the response criteria on the active lever resulted in the delivery of an intravenous nicotine infusion (0.03 ml infusion volume). Responses on the inactive lever were recorded but had no scheduled consequences. Once stable responding at the 0.03 mg/kg/infusion dose was achieved (~5-7 days), animals were then permitted to respond for 0.1 mg/kg/infusion dose of nicotine (training dose) for at least 7 days prior to the start of dose-response assessment. For dose-response studies, animals were permitted to respond for each dose of nicotine for 3-5 days; the mean intake over the last 3 sessions for each dose was calculated and used for statistical analysis. Nicotine doses were presented in ascending order with saline last.

**Electrophysiological recordings**

Adult B6.Cg-Tg(*Chat-COP4*H134R/EYFP*)6Gfng/J mice were sacrificed by cervical dislocation and brains were dissected in chilled (4°C) artificial cerebrospinal fluid (ASCF) containing (in mM): 87 NaCl, 2 KCl, 0.5 CaCl2, 7 MgCl2, 26 NaHCO3, 1.25 NaH2PO4, 25 glucose, 75 sucrose, bubbled with a mixture of 95% O2/5% CO2. 250 µm coronal IPN containing slices were cut with a VT1200S vibratome (Leica, IL, USA), preincubated for 30 min at 37°C and then transferred to the recording solution containing (in mM): 125 NaCl, 2.5 KCl, 2 CaCl2, 1.3 MgCl2, 26 NaHCO3, 1.25 NaH2PO4, 10 glucose, 2 sodium pyruvate, 3 myo-inositol, 0.44 ascorbic acid, bubbled with a mixture of 95% O2/5% CO2. Slices rested in this recording solution at room temperature for at least one hour before they were transferred to the recording chamber. Patch pipettes had resistances of 4–8 MΩ when filled with a solution containing (in mM): 105 K-gluconate, 30 KCl, 10 Hepes, 10 phosphocreatine, 4 ATP-Mg2+, 0.3 GTP (pH adjusted to 7.2 with KOH). Electrophysiological responses were recorded at 33–36 °C with an EPC 10 patch-clamp amplifier and PatchMaster and FitMaster software (HEKA Elektronik, Germany).

Light-induced EPSCs were recorded from IPN neurons of *Chat-ChR2* mice and *Chat-ChR2×Gpr151-KO* mice. Two months old adult animals were used in these experiments. Optic fiber was placed above the IPN region of the acute brain slices. Photo stimulation of IPN cell bodies consisted of a single 5ms pulse of 473nm light delivered at a light power of 5mW from the tip of the optic fiber. Picrotoxin (100 µM) or Forskolin (25 µM) were present to block Gabaergic transmission and activate adenylyl cyclase. For assessments of presynaptic release probability, two paired photo stimuli with an interval of 100ms were delivered. Paired pulse responses of IPN neurons were recorded in the presentce of Nicotine (1µM) and Picrotoxin (100 µM). The paired pulse ratio was defined as the amplitude of the second EPSC to that of the first EPSC.

**Measurement of cAMP levels in dissected IPN brain samples**

IPN samples were accurately dissected from WT and *Gpr151-KO* mice. Tissues were homogenized with a motorized tissue grinder, lysed with 300µl 0.1 mM HCl for 5 min, and centrifuged (14,000 × g) at 4 °C for 10 min. The concentration of cAMP in the supernatant obtained from mouse extracts was measured with the Monoclonal Anti-cAMP Antibody Based Direct cAMP ELISA Kit, Non-acetylated Version (NewEast Biosciences Inc) following the manufacturer’s instructions.

**LANCE cAMP cell-assay measurement**

Lance cAMP measurements were done under forskolin stimulation in HEK 293 and CHO-K1 parental and GPR151-expressin cell lines. When indicated cells were treated with pertussis toxin (PT) (250 ng/ml) overnight. Cells were grown and assayed in culture reagents (DMEM, FBS, BSA) supplied by Life Technologies,Sigma-Aldrich, and Corning plasticware (Corning). Isobutylmethylxanthine (IMBX) and forskolin were obtained from Tocris Bioscience and Sigma-Aldrich. The LANCE cAMP kit and white opaque 384-well plates were purchased from Perkin Elmer. Cells were detached with Versene (Fisher Scientific) and centrifuged to remove supernatant. Then cells were adjusted to 1000 cell/ µL with HBSS stimulus buffer plus Alexa Flour-647 anti-cAMP antibody. 10 µL of cells were dispersed into 384 well plates with MultiDrop Combi. Forskolin was added with HP D300 digital dispenser. Compound was incubated with cells for 45 mins and detection buffer 10 µL was added. The TR-FRET assay (665nM) was further developed for 1 hour and recorded with Envision Lance dual laser 384 protocols.

**SNAP staining on cell lines**

GPR151 was labeled with the cell impermeable SNAP dye 549 from New England Biolabs (NEB). SNAP cell staining was done following the manufacturer’s instructions. Briefly, dye was dissolved in DMSO at 1mM and further diluted into the cell culture medium to a final concentration of 5µM. For imaging, cells were seeded to Matrigel (Corning) coated glass bottom 384-well plate (Aurora) overnight. Then, the medium was removed and 20µL of 5µM dye was added to each well. The samples were incubated at 37ºC for 20 mins. Afterwards, the dye/medium mixture was discarded and cells were washed twice with fresh cell culture medium and once with PBS (with Ca^2+^ and Mg^2+^). Finally, cells were imaged either directly or after being fixed with 2% PFD.
